# Circadian programming of the ellipsoid body sleep homeostat in *Drosophila*

**DOI:** 10.1101/2021.10.22.465404

**Authors:** Tomas Andreani, Clark Rosensweig, Shiju Sisobhan, Emmanuel Ogunlana, William Kath, Ravi Allada

**Affiliations:** Department of Neurobiology, Northwestern University, Evanston, IL, 60208 USA; Department of Engineering Sciences and Applied Mathematics, Northwestern University, Evanston, Illinois 60208

## Abstract

Homeostatic and circadian processes collaborate to appropriately time and consolidate sleep and wake. To understand how these processes are integrated, we scheduled brief sleep deprivation at different times of day in *Drosophila* and find elevated morning rebound compared to evening. These effects depend on discrete morning and evening clock neurons, independent of their roles in circadian locomotor activity. In the R5 ellipsoid body sleep homeostat, we identified elevated morning expression of activity dependent and presynaptic gene expression as well as the presynaptic protein BRUCHPILOT consistent with regulation by clock circuits. These neurons also display elevated calcium levels in response to sleep loss in the morning, but not the evening consistent with the observed time-dependent sleep rebound. These studies reveal the circuit and molecular mechanisms by which discrete circadian clock neurons program a homeostatic sleep center.

## Introduction

The classic two process model posits that the circadian clock and the sleep homeostat independently regulate sleep (Borbely, 1982; Borbely et al., 2016). The circadian process, via phased activity changes in central pacemaker neurons, times and consolidates sleep-wake (Patke et al., 2020). The less well understood homeostatic process, often assayed after extended sleep deprivation, promotes sleep length, depth, and amount as a function of the duration and intensity of prior waking experience (Deboer & Tobler, 2000; Franken et al., 1991; Huber et al., 2004; Werth et al., 1996). Sleep homeostasis is thought to be mediated by the accumulation of various wake-dependent factors, such as synaptic strength (Tononi & Cirelli, 2014), which are subsequently dissipated with sleep.

While homeostatic drive persists in the absence of a functioning circadian clock(Tobler et al., 1983), homeostatic drive can be modulated by the circadian clock. Abolishing clock output through mutation of most core clock genes (Franken et al., 2006; Laposky et al., 2005; Wisor et al., 2002) or electrolytic ablation of the mammalian circadian pacemaker, the suprachiasmatic nuclei (SCN) (Easton et al., 2004) reduces SD-induced changes in non-rapid eye movement (NREM) sleep, an indicator of homeostatic sleep drive in mammals. As circadian clock genes and even the SCN may regulate processes that are not themselves rhythmic(F. Fernandez et al., 2014; McDonald & Rosbash, 2001), these studies leave open the question about whether homeostasis is circadian regulated. To more definitely address the interaction between the clock and the homeostat, sleep-wake have been scheduled to different circadian times in forced desynchrony protocols(Dijk & Czeisler, 1994, 1995). In one such protocol, sleep and wake are scheduled to occur every 28 hours, allowing the circadian clock to free-run with a ∼24 h period. Under these conditions, a variety of indicators of homeostatic drive such as total time asleep, latency to sleep, and NREM sleep time are reduced in the evening independent of time awake (Dijk & Czeisler, 1994, 1995; Dijk & Duffy, 1999; Lazar et al., 2015), consistent with the idea that the clock sustains wakefulness at the end of the waking period in the evening. Yet the molecular and circuit mechanisms by which the circadian clock modulates sleep homeostasis remain unclear.

To understand the mechanistic basis of circadian regulation of sleep homeostasis, we are using *Drosophila,* a well-established model for investigating the molecular and neural basis of circadian rhythms and sleep. Sleep in flies is characterized by quiescence, increased arousal thresholds, changes in neuronal activity, and circadian and homeostatic regulation(Campbell & Tobler, 1984). Flies display each of these hallmarks (Hendricks et al., 2000; Shaw et al., 2000; van Alphen et al., 2013) and have simple, well characterized circadian and sleep neural networks (Dubowy & Sehgal, 2017; Shafer & Keene, 2021). About 150 central pacemaker neurons that express molecular clocks (Dubowy & Sehgal, 2017). Of these, four small ventral lateral neurons (sLNvs) expressing pigment dispersing factor (PDF) are necessary for driving morning activity in anticipation of lights on and exhibit peak levels of calcium around dawn (∼ZT0) (Grima et al., 2004; Liang et al., 2019; Liang et al., 2017; Stoleru et al., 2004). The dorsal lateral neurons (LNds) and a 5th PDF^-^ sLNv are necessary for evening anticipation of lights off and show a corresponding evening calcium peak (ZT8-ZT10) (Grima et al., 2004; Guo et al., 2014#22; Liang et al., 2019; Liang et al., 2017; Stoleru et al., 2004). The posterior DN1 (DN1ps) consist of glutamate-positive (Glu^+^) subsets necessary for morning anticipation and Glu^-^ necessary for evening anticipation under low light conditions (Chatterjee et al., 2018). Lateral posterior neurons (LPN) are not necessary for anticipation but are uniquely sensitive to temperature cycling (Miyasako et al., 2007). Specific pacemaker subsets have been linked to wake promotion (PDF^+^ large LNv(Chung et al., 2009; Parisky et al., 2008; Sheeba et al., 2008), diuretic hormone 31 (DH31^+^) DN1ps(Kunst et al., 2014)) and sleep promotion (Glu^+^ DN1ps (Guo et al., 2016), Allostatin A^+^ LPNs (Ni et al., 2019)), independently of their clock functions. How these neurons regulate homeostatic sleep drive itself remains unsettled.

Timed signaling from these clock neurons converges on the neuropil of the ellipsoid body (EB). The sLNvs and LNds may communicate to R5 EB neurons possibly through an intermediate set of dopaminergic PPM3 neurons based largely on correlated calcium oscillations(Liang et al., 2019). The anterior projecting subset of DN1ps provide sleep promoting input to other EB neurons (R2/R4M) via tubercular bulbar (TuBu) interneurons (Guo et al., 2018; Lamaze et al., 2018). Activation of a subset of these TuBu neurons synchronizes the activity of the R5 neurons which is important for sleep maintenance (Raccuglia et al., 2019). Critically, the R5 neurons are at the core of sleep homeostasis in *Drosophila* (Liu et al., 2016). R5 neuronal activity is both necessary and sufficient for sleep rebound(Liu et al., 2016). Extended sleep deprivation (12-24h) elevates calcium, the critical presynaptic protein BRUCHPILOT (BRP), and action potential firing rates in R5 neurons. The changes in BRP in this region not only reflect increased sleep drive following SD but also KD of *brp* in R5 decreases rebound (Huang et al., 2020) suggesting it functions directly in sleep homeostasis. R5 neurons stimulate downstream neurons in the dorsal fan-shaped body (dFB), which are sufficient to produce sleep (Donlea et al., 2014; Donlea et al., 2011; Liu et al., 2016). Yet how the activity of key clock neurons are integrated with signals from the R5 homeostat to determine sleep drive remains unclear.

Here we dissect the link between the circadian and homeostatic drives by examining which clock neural circuits regulate sleep rebound at different times of day in *Drosophila*. Akin to the forced desynchrony protocols, we enforced wakefulness at different times of day and assessed sleep rebound. We exposed flies to 7 h cycles of sleep deprivation and recovery, enabling assessment of homeostasis at every hour of the day. We found that rebound is suppressed in the evening in a *Clk*-dependent manner. We demonstrate that these effects are mediated by specific Glu^+^ DN1p pacemaker neurons in the morning and PDF^-^ LNd/sLNv in the evening, independent of their effects on locomotor activity. Moreover, homeostatic R5 EB neurons integrate circadian timing and homeostatic drive; we demonstrate that activity dependent and presynaptic gene expression, BRP expression, neuronal output, and wake sensitive calcium levels are all elevated in the morning compared to the evening, providing an underlying mechanism for clock programming of time-of-day dependent homeostasis.

## Results

### Scheduled sleep deprivation demonstrates suppression of rebound in the evening

To confirm and resolve the timing of clock modulation of sleep rebound, we scheduled sleep deprivation in flies at different times of day and assessed sleep rebound, a protocol we term scheduled sleep deprivation (SSD). We employed an ultradian 7h cycle over 7 days allowing us to observe rebound at each hour of the 24 hour day (24 total deprivations) (Fig. 1a,b). SD was administered for 2.5 hours followed by 4.5 hours of rebound such that flies would be allowed ∼⅔ of the day to sleep, similar to the ratio of sleep observed in a WT female fly without SD. Given the potential for stress effects of longer deprivation typically used in flies (6-24h) we opted for a shorter 2.5 h protocol. Indeed, there was no significant difference between total sleep in flies kept in SSD and those under baseline conditions (Fig. 1d). In addition, sleep rebound does not increase over the course of the 7 day protocol further suggesting that flies are able to fully recover sleep during the 4.5 h rebound period (Fig. 1e). To test if SSD modulated the circadian phase, SSD flies released into constant dark (DD) following the protocol did not exhibit any detectable change in phase (Fig. 1c). Together these results demonstrate that the SSD protocol allows assessment of rebound at different times of day without altering total sleep or circadian phase.

**Figure 1:**
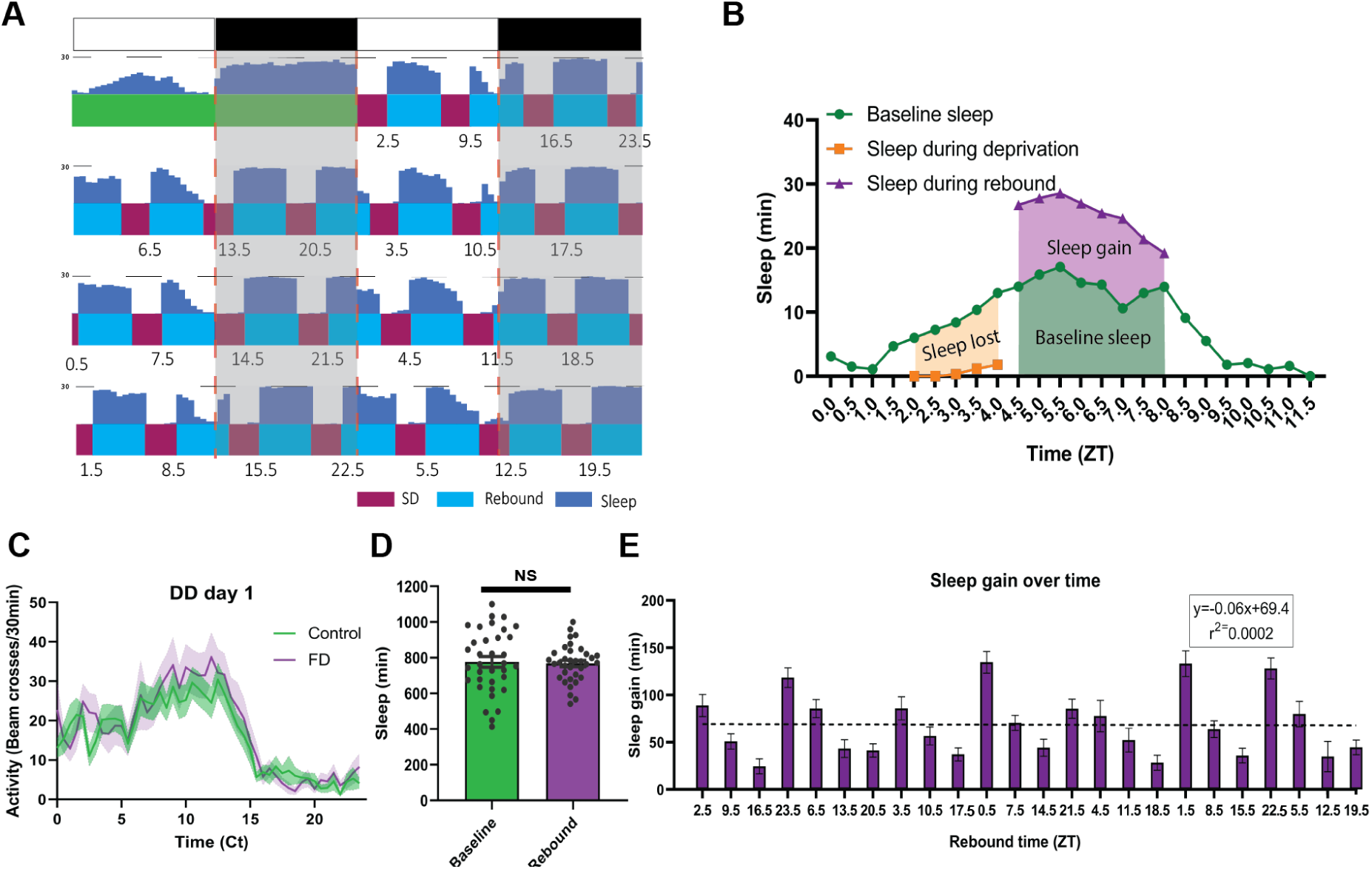
T7 Drosophila forced desynchrony protocol can be used to illustrate time dependent rebound. (A) Average WT sleep (N=32) over the final 8 days of FD protocol with the time at which rebound begins (ZT) noted below each rebound period. (B) Profiles of sleep metrics used to compare rebound at different times of day (example is rebound occurring at ZT4.5). Sleep lost is determined by the difference between baseline sleep and sleep during the SD. Sleep gain is determined by the difference between rebound and baseline sleep. (C) Average activity of WT flies over 24 hours of flies released into the dark following FD stimulation (N=19) or control (N=19) that received no stimulation. WT Flies released into DD1 following FD display a profile of activity similar to control flies. Shaded bands indicate SEM.(D) Average sleep during baseline and the average sleep per day during the 7 day SD-rebound period (individual flies shown circles). There is no significant difference between average baseline sleep and average sleep per day over the course of the FD (P>0.08, paired t-test). (E) Average WT (N=32) sleep gain across the course of the experiment with rebound time(ZT) depicted on the x axis. Regression of WT sleep gain over the course of the experiment displays no significant trend (P>0.95 linear regression). Data are means +∕- SEM.

By comparing flies’ baseline sleep to their rebound sleep (sleep after deprivation) around the clock, we observed robust rebound in the morning and suppressed rebound in the evening (Fig. 2a). Under baseline conditions, flies typically show morning and evening peaks in wakefulness/activity. After sleep deprivation, flies display a robust sleep rebound throughout the 4.5 h rebound period in the morning while evening rebound is suppressed (Fig. 2a). To statistically compare morning and evening times of day here and throughout this study, we selected specific time points where the amount of sleep deprived and the baseline sleep during the rebound, two potential confounds, were comparable, allowing a direct comparison of sleep rebound. As indicated in the heat map, we found sleep rebound in the morning is significantly higher than sleep rebound in the evening when controlling for baseline sleep such that there is a >2x difference in rebound between morning and evening time points (rebound at ZT1.5∼133 min and ZT9.5∼51 min) (Fig. 2c). This was also accompanied by a significant difference in latency following deprivation (Supplemental Fig. 2c). We observed similar results using a streamlined protocol focusing on morning (ZT1.5 and 2.5) and evening timepoints (ZT8.5, 9.5, 10.5) (Supplemental Fig 1). During the course of our experiments, we transitioned to a more streamlined protocol to reduce the length of the protocol and the number of sleep deprivations, minimizing the potential for trends in sleep over the course of the protocol. Video evidence confirms that these morning/evening differences are not due to failure to cross the infrared beam due to increased feeding (Supplemental Videos 1,2). Lastly, we determined if these effects persist under constant darkness (DD). We observed elevated rebound in the morning (CT2.5) relative to the evening (CT10.5), indicating that these differences are not dependent on light (Fig. 2e). All together, this data suggests that homeostatic rebound sleep is strongly modulated by the internal clock.

**Figure 2:**
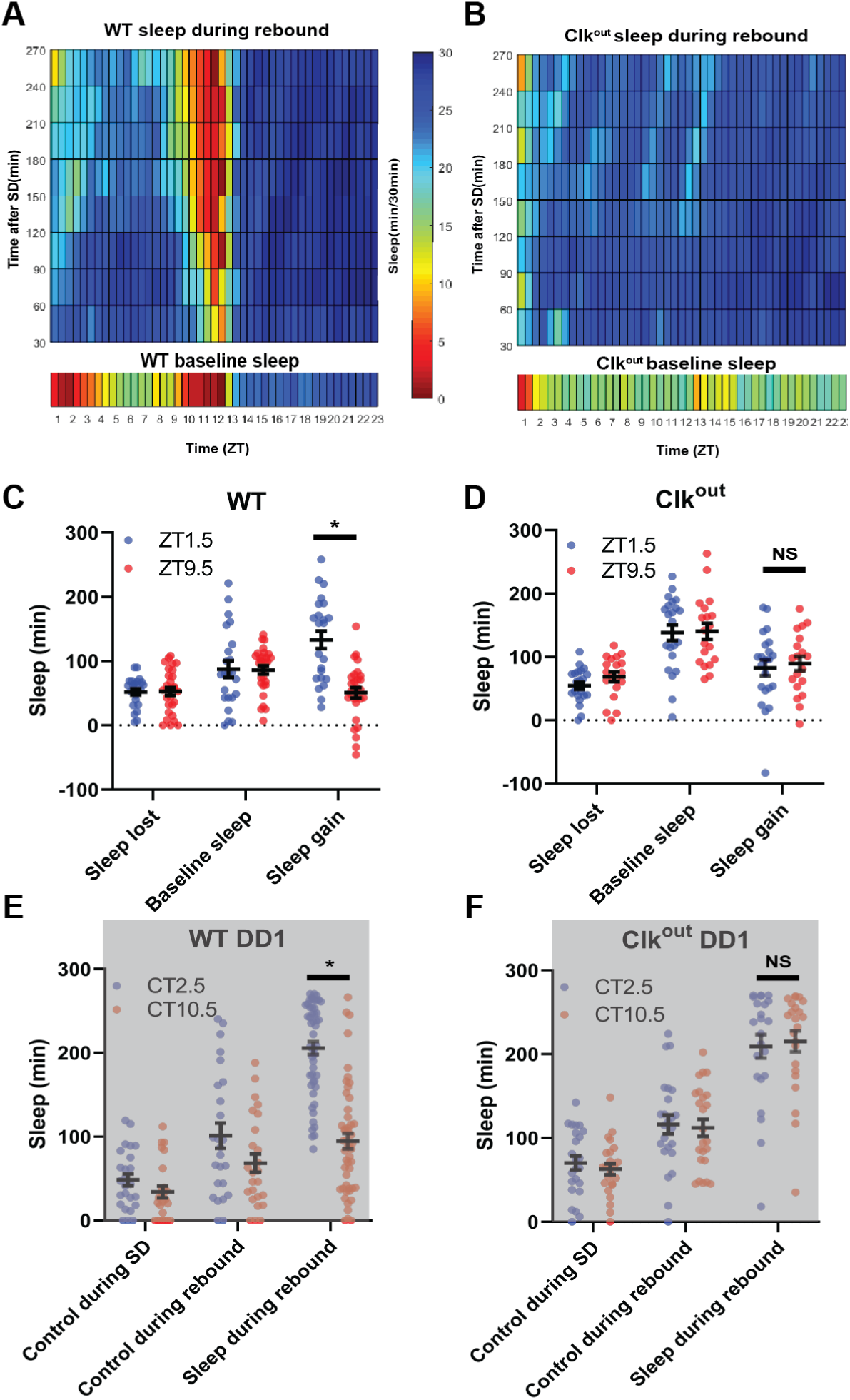
Sleep rebound is dependent on the molecular clock. (A-B) Rebound sleep heatmaps (above) illustrate average sleep as a function of time of day when rebound occurred (ZT) and minutes after SSD episode. Missing time points are filled using matlab linear interpolation function. Baseline sleep heatmaps (below) illustrate average sleep during 30 min bins. (A) WT (N=32) baseline displays low sleep following lights on and preceding lights off. Immediately following SD flies show high sleep except in the hours preceding lights off. Flies tend to sleep less as rebound time proceeds. (B) Clk^out^ (N=40) baseline sleep (below) is nearly constant except for low sleep immediately following lights on. SD uniformly increases sleep and flies tend to sleep less as rebound time proceeds. (C-D) Comparison of sleep lost, baseline sleep, and sleep gain following deprivation at morning and evening timepoints in WT and clk^ou,^. (C) Sleep gain is greater for WT (N=32) rebound at ZT1.5 compared to ZT9.5 (P<.00001, paired t-test). (D) No difference between sleep gain at the two time points is observed in Clk^ou,^ (N=40) (P>0.37, paired t-test). (E,F) Two sleep measures in control flies (control during SD and control during rebound), along with sleep during rebound in SD with rebound at 2.5 and 10.5. (F) Rebound sleep is greater following deprivation at CT2.5 compared to CT10.5 (P<.00001, paired t-test) in WT flies (N= 49). (G) No difference in rebound sleep is observed in Clk^ou,^ (N= 23) (P>0.75, paired t-test). Data are means +∕- SEM

### Sleep rebound is dependent on the molecular clock

To determine if morning/evening differences in rebound are due to the circadian clock we performed SSD in arrhythmic *Clk*^out^ (Lee et al., 2014) and short-period *per^s^* mutants, which have an advanced evening peak in LD (Hamblencoyle et al., 1992; Konopka & Benzer, 1971). In the absence of *Clk*, flies do not display the wild-type morning and evening peaks of wakefulness and exhibit robust rebound at all times, reaching maximal levels of sleep after each SD (Fig. 2b). Selected morning/evening time points do not exhibit significant differences in rebound in LD (ZT1.5 and ZT8.5) nor in DD (ZT2.5 and ZT10.5) (Fig. 2d, f). There was also no difference in latency between matched morning and evening time points (ZT1.5 and ZT8.5) after sleep deprivation in *Clk*^out^ (Supplemental Fig. 2d). Similar to wild-type flies, *per^s^* showed elevated rebound in the morning compared to the evening; however, as expected, the trough of rebound sleep in the evening was phase advanced relative to wild-type by about 4 hours (ZT5.5 v. ZT9.5) (Supplemental Fig. 2a, b). Furthermore, *per^s^* flies exhibit an increased sleep latency following deprivation in earlier evening time points (ZT7.5) relative to control (ZT9.5) (Supplemental Fig. 2e). The loss of a morning/evening difference in rebound in arrhythmic *Clk*^out^ and the phase advance of evening rebound suppression in *per^s^* further support the role of the clock in regulating sleep rebound.

### Glutamatergic DN1p circadian pacemaker neurons mediate morning and evening differences in rebound

To address the underlying neuronal basis, we employed a “loss-of-function” approach where we inactivate and/or ablate targeted neuronal populations and assess the impact on sleep rebound at different times of day. To test the role of clock neurons, we selectively ablated subsets by expressing the pro-apoptotic gene *head involution defective* (*hid*) using the Gal4/UAS system. Ablation of most of the pacemaker neurons including those underlying morning and evening behavior using *cry39-Gal4* (Klarsfeld et al., 2004; Picot et al., 2007) substantially reduced both morning and evening anticipation in males (Supplemental Table 1) as previously described (Grima et al., 2004). Anticipation in females is more difficult to quantify due to more consolidated sleep and wake, i.e., sleep at night reduces morning anticipation, more mid-day wake reduces evening anticipation(Isaac et al., 2010). Consistent with the loss of circadian function, ablation also abolished the difference between morning and evening rebound (Supplemental Fig. 3a, b), predominantly by elevating evening rebound (at ZT9.5 control ∼27 min, *cry39-Gal4* ∼100 min). We ablated PDF+ using *pdf-Gal4*, and despite substantially reduced morning anticipation in males validating our reagent (Supplemental table 1), the morning/evening difference in rebound nonetheless persists when comparing morning/evening time points (Supplemental Fig. 3c). Coupling *cry39-Gal4* with *pdf-Gal80* to ablate most clock cells except PDF^+^ neurons confirms this observation; these flies display comparably high rebound between morning and evening time points similar to *cry39-Gal4* (Supplemental Fig. 3d), highlighting the role of non-PDF clock neurons.

A potential synaptic target of the PDF^+^ sLNv that are also important for morning behavior are the Glu^+^ DN1p neurons(Chatterjee et al., 2018; L. Zhang et al., 2010; Y. Zhang et al., 2010). Targeting of the Glu^+^ DN1p has relied on drivers that are expressed outside of the DN1p including other sleep regulatory neurons(Chatterjee et al., 2018; Guo et al., 2016). To more definitively test their function, we employed the intersectional split Gal4 system (Dionne et al., 2018) utilizing two promoters, *R18H11* (expressed in DN1p and other neurons)(Guo et al., 2016) and *R51H05* that uses the vesicular glutamate transporter (vGlut) promoter presumably targeting glutamatergic neurons. This intersection resulted in expression in just 6-7 neurons per hemisphere with little or no expression elsewhere in the brain (Fig. 3a, b). We targeted *hid* expression using this split Gal4, we observed a reduction in morning anticipation in males demonstrating the necessity of this defined neuronal group (Supplemental Table 1). However, in females used in our protocols, we did not observe a reduction in morning anticipation, possibly due to the lights-on activity peak masking anticipation (Fig. 3 e, f). We also did not observe significant changes in baseline sleep levels (Fig. 3g). Despite the lack of a significant change in their baseline sleep/activity profiles, ablation eliminated the difference in morning and evening rebound (Fig. 3c, d). This change appears to be primarily due to a reduction of morning rebound (at ZT1.5 control ∼104 min, Glu^+^ DN1p ∼59 min). Thus, using highly specific drivers, we find that Glu+ DN1ps promote rebound sleep in the morning largely independent of their role in regulating baseline sleep/activity.

**Figure 3:**
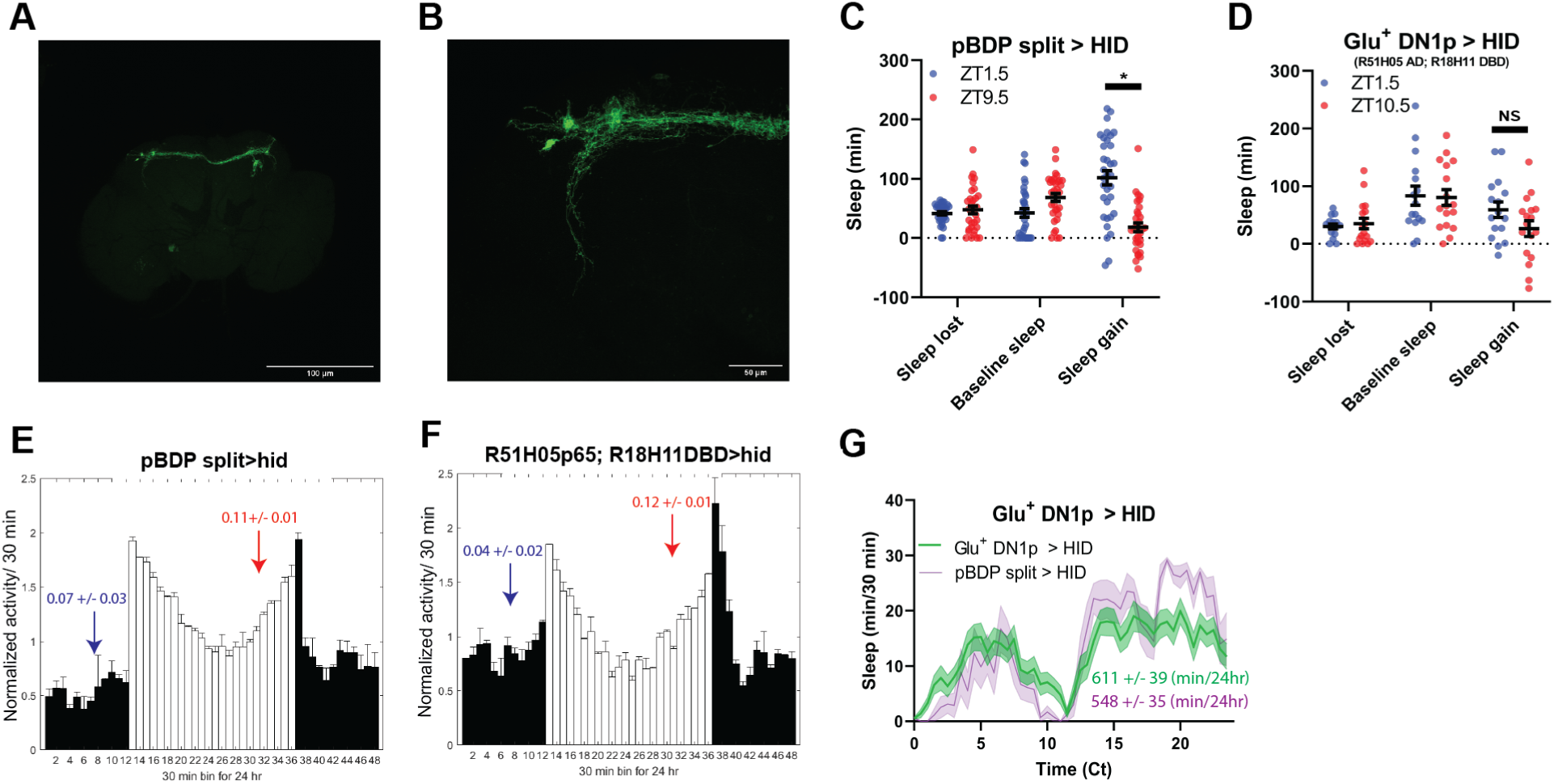
Glutamatergic DNlps enhance morning rebound. (A-B) GFP Expression pattern of split Gal4 line that labels Glu+ Dnlps (R51HO5 AD; R18H11 DBD > GFP) at 10x (A) and 40x (B). (C-D) Comparison of sleep lost, baseline sleep, and sleep gain following deprivation at morning and evening timepoints in glutamatergic DNlp ablated flies. Morning times are matched with evening time points with similar baselines. (C) Control flies with no ablated neurons (pBDP split > hid) (N=26) exhibit greater rebound in the morning compared to matched evening time point (P<0.0001, paired t-test). (D) Flies with Glu+ DNlps ablated (R51H05 AD; R18H11 DBD > hid) (N=14) do not exhibit a significant difference in sleep gain between matched moming/evening time points (P>0.09, paired t-test). Data are means +∕- SEM. (E-F) Averaged activity eductions for female flies during the first 2 days of 12:12 LD. The light-phase is indicated by white bars while the dark-phase is indicated by black bars. Morning and evening anticipation indices are represented in blue and red respectively.(G) Average sleep during the baseline day. Glu+ DNlps ablated (R51HO5 AD; R18H11 DBD > hid) (N=30) (green) and control (pBDP split > hid) (N=26) (purple). Sleep per 24 hours is indicated in the bottom right. Data are means +∕- SEM.

### TuBu and R2/R4m neurons are important for time-dependent modulation of sleep homeostasis

A subset of DN1ps send anterior projections to TuBu interneurons which in turn target the R2/R4m neurons of the EB(Guo et al., 2018; Lamaze et al., 2018)(Fig. 4a). TuBu neurons are a heterogeneous group distinguished by their axonal projections to 3 regions (superior, anterior and inferior) of the Bulb (BU), a neuropil comprised of, among other things, dendritic projections of neurons that form the EB (Lovick et al., 2017; Omoto et al., 2017). Previous studies have highlighted the role of the superior projecting TuBu neurons in generating sleep (Guo et al., 2018; Lamaze et al., 2018). To validate and further resolve this circuitry, we mined the Janelia Farm connectome which uses a large-scale reconstruction of the central brain from electron microscopy data(Scheffer et al., 2020). Using this approach, we identified direct synaptic connections from a subset of DN1pB (body IDs: 386834269, 5813071319) to a subset of TuBu neurons (TuBu01), to R4m neurons and eventually to R2 neurons (Supplemental Fig. 4a,b). Based on their morphology the Tubu01 neurons are anterior\inferior projecting. Thus, this connectome analysis both validated this circuit but also provided higher resolution for specific subsets that may be involved.

**Figure 4:**
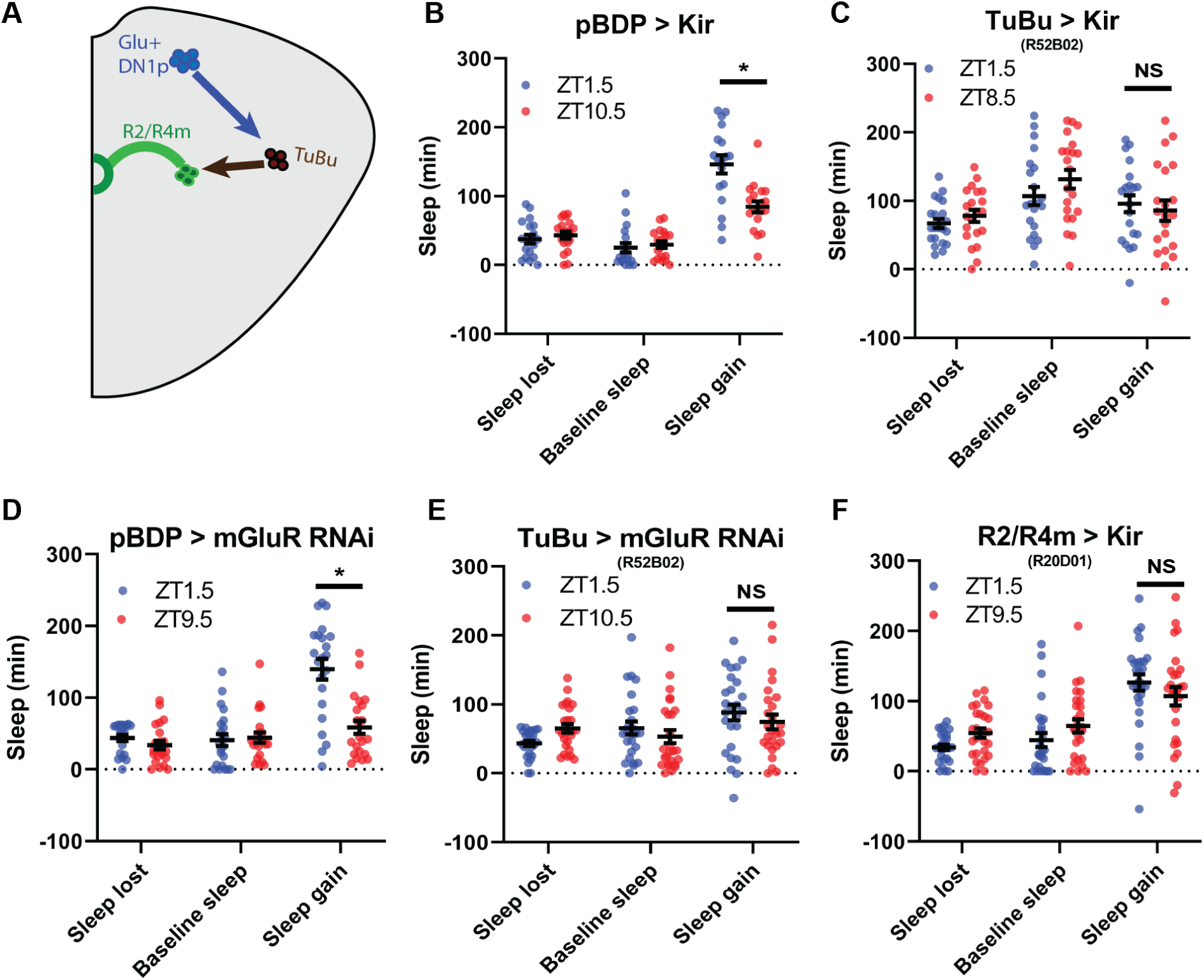
TuBu intermediates convey enhanced morning glutamatergic signal to R2∕R4m ellipsoid body neurons. (A) Cartoon illustrating proposed link between Glu+ DNlps and R2∕R4m with Tubu intermediates. (B-F) Comparison of sleep lost, baseline sleep, and sleep gain following deprivation at morning and evening timepoints while modulating neurons linking DNlps to the EB. Morning times are matched with evening time points with similar baselines. (B) Enhancerless-Gal4 control flies (pBDP > Kir) (N=21) exhibit greater rebound in the morning compared to a matched evening time point (P<0.01, paired t-test). (C) Flies with TuBu neurons silenced (R52B02 > Kir) (N=21) do not exhibit a difference in rebound between matched moming/evening time points(P> 0.38, paired t-test). (D) Enhancerless-Gal4 driver paired with UAS-GluR-RNAi (pBDP > GluR RNAi) control (N=32) exhibit greater rebound in the morning compared to matched evening time point (P<0.00001, paired t-test). (E) Flies with KD of GluR in TuBu neurons (R52B02 > GluR RNAi) do not exhibit a significant difference between matched moming/evening time points (P>0.28, paired t-test). (F) Flies with R2∕R4m neurons silenced (R20D01 > Kir) (N=32) do not exhibit a significant difference in rebound between matched moming/evening time points(P>0.26, paired t-test). Data are means +∕- SEM.

To determine if these neurons are important for sleep homeostasis, we first tested Gal4 drivers previously used to mark these neurons (Guo et al., 2018; Lamaze et al., 2018; Liang et al., 2019; Liu et al., 2016) in combination with *hid*, but found that in many cases (*R52B02*, *R20D01*) they were lethal, likely due to broader anatomic and/or developmental expression. So instead we used the inward rectifying potassium channel Kir2.1(Baines et al., 2001) to silence these neurons and examined sleep rebound in the morning and evening. Silencing of a previously used driver (*R92H07*) that labels superior projecting TuBu neurons had no effect on rebound (Supplemental Fig. 4c,d). We identified another GAL4 driver (*R52B02*) that labels the superior and anterior and/or inferior subgroups previously implicated in sleep regulation(Guo et al., 2018; Jenett et al., 2012; Lamaze et al., 2018; Pfeiffer et al., 2008). We used this line in combination with Kir2.1 and found that the difference between morning and evening rebound was lost, similar to what was observed after Glu^+^ DN1p ablation (Fig. 4b,c). We knocked down the expression of a metabotropic glutamate receptor (mGluR) in these neurons using RNAi (Guo et al., 2016) and observed phenotypes very similar to silencing them (Fig 4d,e). To determine which neurons are acting downstream of TuBu, we targeted the R2/R4m neurons using *R20D01*(Lamaze et al., 2018). Silencing these neurons with Kir2.1 eliminated the difference between rebound in the morning and evening, phenocopying Glu^+^ DN1p ablation and TuBu silencing (Fig. 4f). Taken together, these results demonstrate a role for the DN1p-Tubu-R2/R4m circuit in regulating time-dependent sleep rebound.

### PDF^-^ sLNv and LNds mediate evening suppression of sleep rebound

To determine the cellular basis of the evening rebound phenotype, we selectively ablated 2-3 LNds and the 5th sLN_v_ (4 neurons) using the highly specific MB122B split Gal4 line(Guo et al., 2017). This manipulation resulted in a large (>6 fold) increase in rebound in the evening (at ZT9.5 control ∼22 min, *MB122B*∼ 141 min) and a more modest (∼1.5 fold) effect in the morning (at ZT1.5 control∼ 104 min, *MB122B* ∼157 min) (Fig. 5a, b). We observed similar results with Kir2.1 (Fig. 5f, g). Surprisingly we did not observe significant effects on baseline sleep levels (Fig. 5c, h) or anticipation by ablation or silencing (Fig. 5d-e, i-j). Differences between these baseline anticipation results and previously observed silencing effects on sleep may be due the use of constitutive versus inducible silencing(Guo et al., 2017). Nonetheless, these results indicate that effects on rebound are largely independent of baseline anticipation/sleep levels. Thus, just 4 PDF^-^ LNd/sLNv cells are essential for clock control of rebound with an especially strong suppressive effect in the evening.

**Figure 5:**
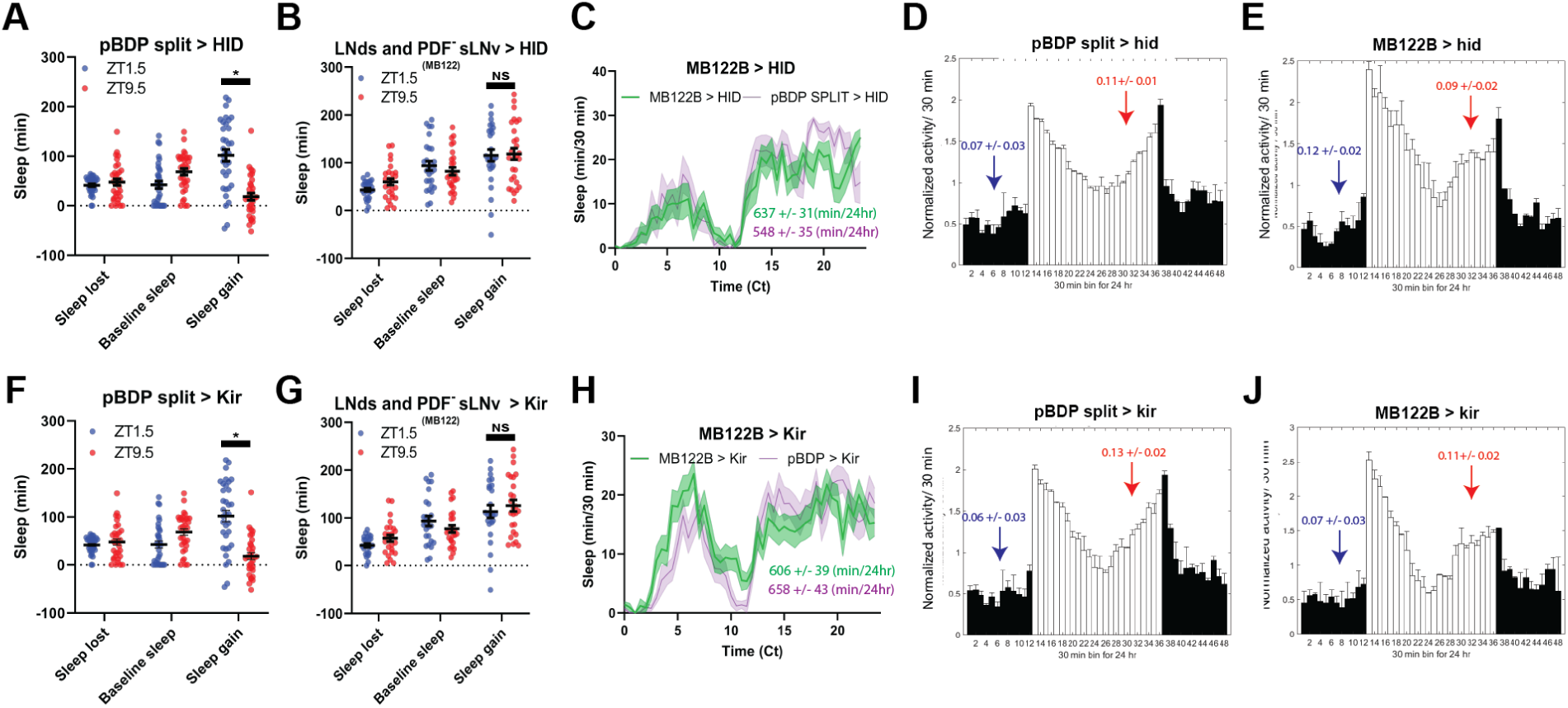
LNds and the PDF-sLNv suppress evening rebound. (A,B,F,G) Comparison of sleep lost, baseline sleep, and sleep gain following deprivation at morning and evening timepoints in clock neuron-ablated flies. Morning times are matched with evening time points with similar baselines. (A) Control flies with no ablated neurons (pBDP split > hid) (N=26) exhibit greater rebound in the morning compared to matched evening time point (P<0.0001, paired t-test). (B) Flies with 2-3 LNds and the PDF- sLNv ablated (MB122B > hid) (N=30) do not exhibit a significant difference in sleep gain between matched moming/eve-ning time points (P>0.50, paired t-test). (F) Control flies with no silenced neurons (pBDP split > kir) (N=34) exhibit greater rebound in the morning compared to matched evening time point (P<0.0001, paired t-test). (G) Flies with 2-3 LNds and the PDF- sLNv silenced (MB122B > kir) (N=31) do not exhibit a significant difference in sleep gain between matched moming/evening time points (P>0.45, paired t-test). Data are means +∕- SEM. (C,H) Average sleep during the baseline day. (C) LNds and the PDF- sLNv ablated (MB122B > hid) (N=30) (green) and control (pBDP split > hid) (N=26) (purple). (H) 2-3 LNds and the PDF- sLNv silenced (MB122B >kir) (N=31) (green) and control (pBDP split > kir) (N=34) (purple).Sleep per 24 hours is indicated in the bottom right. (D,E,I,J) Averaged activity eductions for female flies during the first 2 days of 12:12 LD. Light phase is indicated by white bars while the dark phase is indicated by black bars. Morning and evening anticipation indices are represented in blue and red respectively difference between morning and evening time points (P>0.45, paired t-test).

### PPM3, R5 and dFB neuron synaptic output is required for intact sleep homeostasis

The PPM3 and R5 neurons have been implicated as downstream of the LNd (Fig 6a). To test the effects of PPM3 on sleep homeostasis we blocked synaptic transmission by expressing tetanus toxin (TNT) (Sweeney et al., 1995) using *R92G05*-*Gal4* (Liang et al., 2019). As LNd calcium oscillations are synchronized with those in the PPM3, we hypothesized that PPM3 silencing may phenocopy LNd ablation, i.e., increasing rebound in the evening. However, PPM3 silencing dramatically reduced rebound in both the morning and the evening with little difference between the two times (ZT1.5 and ZT8.5), suggesting that PPM3 are not mediating LNd effects (Fig. 6c,d). Like the PPM3 neurons, blocking R5 synaptic output using a novel split GAL4 (*R58H05 AD; R48H04 DBD*) (Fig 6b) also reduced rebound in both morning and evening, consistent with the role of these neurons in mediating rebound from 12 h SD (Liu et al., 2016) (Fig 6e,f). Moreover, no difference between morning and evening rebound was evident. R5 neurons promote sleep in response to deprivation by activating the sleep promoting dFB (Liu et al., 2016). Thus, we also blocked synaptic output from the dFB using TNT. Rebound was reduced as previously reported(Qian et al., 2017) but without any morning/evening difference, just as it was for PPM3 and R5 (Fig. 6g,h). Although the exact nature of the PPM3 input remains an open question, these studies highlight a role for a PPM3-R5-dFB pathway in rebound sleep in response to deprivation at all times of day even with shorter deprivation protocols.

**Figure 6:**
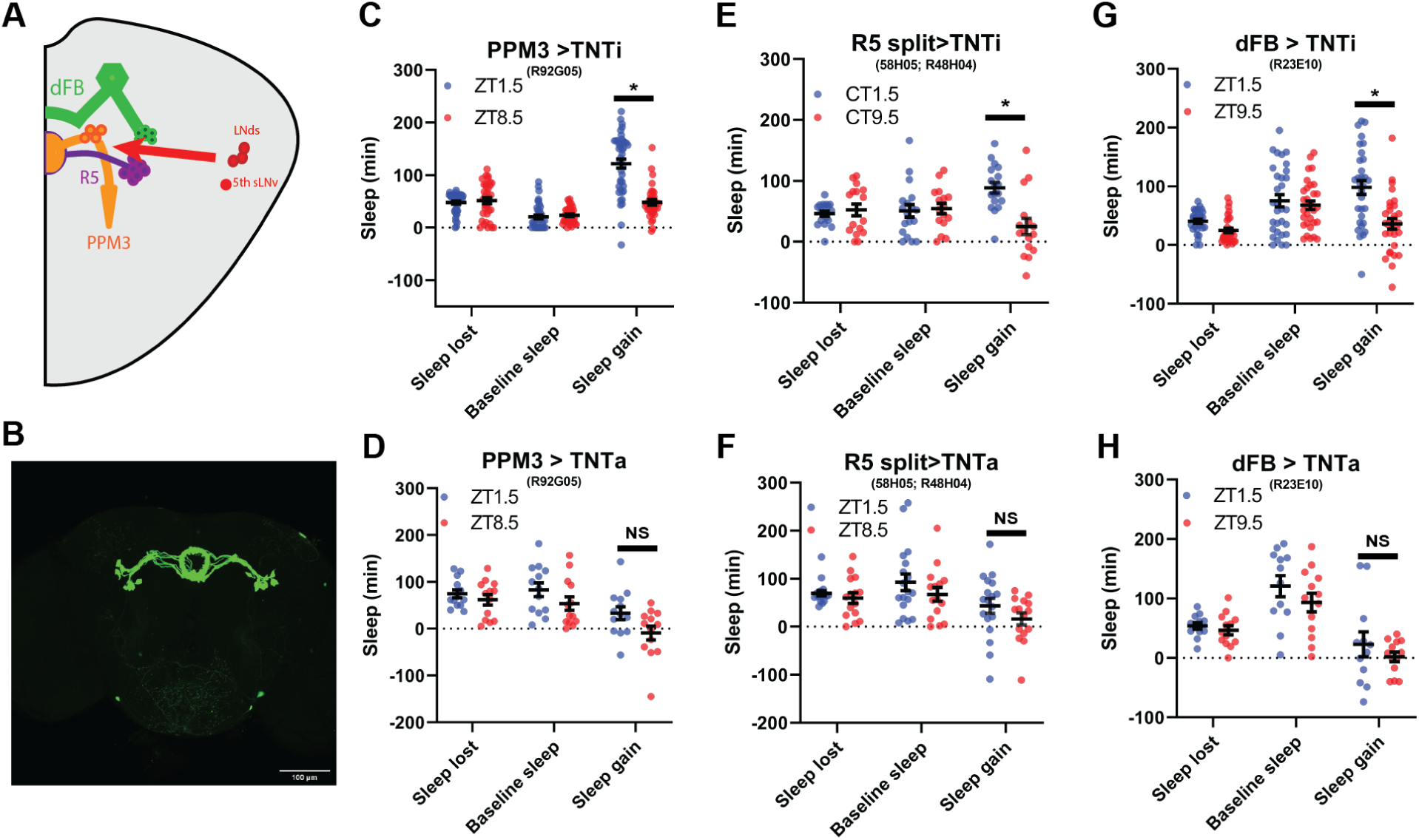
PPM3 supply evening suppressing homeostatic signal to R5 ellipsoid body neurons. (A) Cartoon illustrating link between LNds and 5th sLNv and dFB via with PPM3 and R5 intermediates. (B) GFP Expression pattern of split Gal4 line that labels Glu+ Dnlps (R58H05 AD; R48H04 DBD > GFP) at 20x. (C-H) Comparison of sleep lost, baseline sleep, and sleep gain following deprivation at morning and evening tιmepomts modulating neurons linking LNd activity to the EB. Morning times are matched with evening time points with similar baselines. (C) Flies expressing an inactive fonn of tetanus toxin in PPM3 neurons (R92G05 > TNTι)(N=45) exhibit greater rebound in the morning than at a matched evening time point (P<0.0001, paired t-test). (D) Silencing PPM3 neurons with an active fonn of tetanus toxin (R92G05 > TNTa)(N=27) resulted in no significant difference between matched mornnιg∕evenmg time points (P>0.10, paired t-test). (E) Flies expressing an inactive fonn of tetanus toxin in R5 neurons (R58H05 AD; R48H04 DBD > TNTi) (N=21) exhibit greater rebound in the morning than at a matched evening time point (P<0.01, paired t-test). (F) Silencing R5 neurons with tetanus toxin (R58H05 AD; R48H04 DBD > TNTa) (N=16) resulted in no significant difference in sleep gam for matched morning and evening time points (P>0.70, paired t-test). (G) Flies expressing an inactive fonn of tetanus toxin in the dFB (R23E10 > TNTi) (N=30) exhibit greater rebound in the morning than at a matched evening time point (P<0.0001, paired t-test). (H) Silencing dFB neurons with tetanus toxin (R23E10 > TNTa)(N=12) resulted in no significant difference between morning and evening time points (P>0.45, paired t-test).

### R5 ellipsoid body neurons exhibit elevated expression of activity-dependent and presynaptic genes in the morning relative to the evening

To ascertain how the circadian system may impact the R5 homeostat, we examined molecular and physiological changes in R5 as a function of time and sleep need. Interestingly, activation and deprivation studies have focused exclusively on morning rebound. To identify time- and wake-dependent gene expression in an unbiased manner, we selectively labeled R5 neurons (Fig 6b, *R58H05 AD; R48H04 DBD > GFP*) and subjected flies to 2.5 h of mechanical SD in either the morning or evening. We then isolated R5 neurons from control or SD flies at ZT1 and ZT9 using fluorescence-activated cell sorting and subjected them to RNA-sequencing.

Based on our behavioral data, we hypothesized that morning SD would induce differential gene expression compared to control flies that did not receive SD while evening SD would not be sufficient to induce changes in gene expression compared to controls. We were surprised to find that neither morning nor evening SD had much of an effect on gene expression in the R5 neurons (Fig 7a,b). In the morning, only two genes were significantly differentially expressed (q<0.1, *Hsp70Bb* and *stv*). Likewise, in the evening, only four genes were significantly differentially expressed (q<0.1, *CG5522, CG13285, mt:ND5,* and *Hsp70Bb*). In stark contrast, comparisons of morning and evening timepoints with or without sleep deprivation (Morning Control (MC) vs Evening Control (EC), Morning SD (MSD) vs Evening SD (ESD), or MC + MSD vs EC + ESD) produces 46-128 differentially expressed genes (q<0.1, Fig 7c,d,e). Notably, this time of day dependent regulation does not appear to be driven by core clock genes in these neurons (Supplemental Fig 5). *Clk* is detected in only 2 out of 12 samples and only at very low levels in those samples. Also the expression of other clock genes like *per* and *tim* is not fluctuating between the two timepoints.

**Figure 7:**
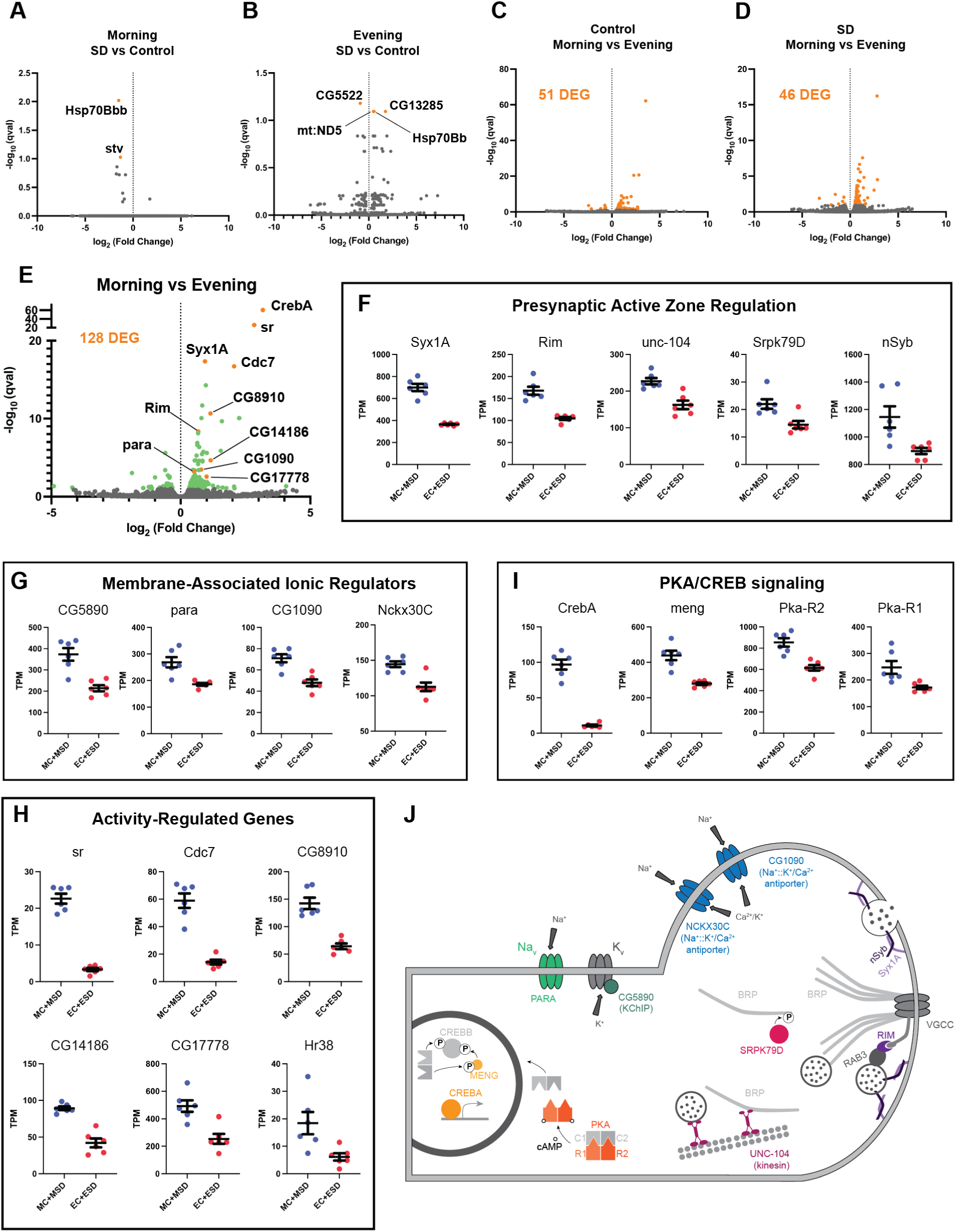
RNA sequencing of FAC-sorted R5 neurons suggests elevated activity in the morning. (A) Volcano plot (fold change versus qval) of Morning SD (MSD) vs Morning Control (MC) gene expression. Significantly differentially expressed genes shown in orange. (B) Volcano plot of Evening SD (ESD) vs Evening Control (MC) gene expression. Significantly differentially expressed genes shown in orange. (C) Volcano plot of MC vs EC gene expression. 51 significantly differentially expressed genes (DEG) were identified and are shown in orange. (D) Volcano plot of MSD vs ESD gene expression. 46 significantly differentially expressed genes (DEG) were identified and are shown in orange. (E) Volcano plot of MC+MSD vs EC+ESD gene expression. Differentially expressed genes are shown in green with a few genes highlighted in orange and labeled. (F-I) Scatter plots for several differentially expressed genes. Transcripts Per Kilobase Million (TPM) is shown for each sample. All morning samples are grouped and all evening samples are grouped. Graphs are grouped by similar functions: (F) active zone components/regulators, (G) membrane-associated ionic regulators, (H) activity-regulated genes, (I) PKA/CREB signaling. (J) Schematic of select morning upregulated genes. Upregulated genes are shown in color while other interacting components are depicted in gray. PARA and CG5890 are both involved in the generation and propagation of action potentials. Multiple active zone components/regulators (NSYB, SYX1A, RIM, SRPK79D, UNC-104) interact with BRP and voltage-gated calcium channels (VGCCs) to support neuronal output and intracellular calcium influx. Elevated levels of intracellular calcium are regulated by the antiporters NCKX30C and CG1090. Second messenger cAMP interacts with regulatory subunits of PKA (R1/R2) and releases the catalytic subunits (C1/C2) to phosphorylate CREBB and MENG, stabilizing CREBB. CREBA acts as a transcriptional activator independent of PKA activity.

To understand what sorts of molecular programs are undergoing differential regulation between morning and evening, we examined gene ontologies of genes upregulated in the morning. These terms include cellular components like “presynaptic active zone”, “synaptic vesicle”, “terminal bouton”, and “cAMP-dependent protein kinase complex”, as well as molecular functions like “calcium ion binding” and “calcium, potassium::sodium antiporter activity”. The genes identified in these categories suggest a temporally regulated state of activity for the R5 neurons. Indeed, major active zone regulators such as *Syx1A*, *Rim*, *unc-104*, *Srpk79D*, and *nSyb* are all significantly upregulated in the morning (Fig 7e,f). *Syx1A*, *Rim*, and *nSyb* are part of the synaptic vesicle docking and exocytosis machinery and *Rim* also regulates the readily-releasable pool of synaptic vesicles, playing a major role in presynaptic homeostasis(Broadie et al., 1995; Muller et al., 2012). *unc-104* is involved in trafficking of synaptic vesicles and BRP to the active zone(Zhang et al., 2017) and the kinase *Srpk79D* regulates trafficking and deposition of BRP at active zones via phosphorylation of its N-terminus(Johnson et al., 2009; Nieratschker et al., 2009). We also observed significant upregulation of genes involved in ionic transport across the plasma membrane, including *para*, a voltage-gated sodium channel (Catterall, 2000; Loughney et al., 1989), and *CG5890*, a predicted potassium channel-interacting protein (KChIP) (Fig 7e,g). Mammalian KChIPs have been shown to interact with voltage-gated potassium channels, increasing current density and conductance and slowing inactivation(An et al., 2000). Two sodium::potassium/calcium antiporters, *CG1090* and *Nckx30C*, were also upregulated (Fig 7e,g). These antiporters function primarily in calcium homeostasis by using extracellular sodium and intracellular potassium gradients to pump intracellular calcium out of the cell when calcium levels are elevated(Haug-Collet et al., 1999). Amongst the most significantly upregulated genes in our dataset, we found six genes that were previously identified as activity-regulated genes in *Drosophila* (ARGs; *sr, Cdc7* (also known as *l(1)G0148*)*, CG8910, CG14186, CG17778, hr38*) (Fig 7e,h). These genes are analogous to immediate early genes in mammals and represent half of a group of twelve genes that were induced in three distinct paradigms of neuronal stimulation(Chen et al., 2016). Finally, we found that several critical components of Creb signaling were enriched in the morning in R5 neurons (Fig 7e,i). *CrebA* was the most significantly upregulated gene in the morning samples, though we also saw significant increases in *meng*, which encodes a kinase that works synergistically with the catalytic subunits of PKA to phosphorylate and stabilize CREBB(Lee et al., 2018), as well as both regulatory subunits of PKA (*Pka-R1, Pka-R2*) (Fig 7e,i). CREBA and CREBB likely serve different roles, but appear to be involved in activity-dependent processes like dendritogenesis and long term memory(Iyer et al., 2013; Yin et al., 1995).

Synthesizing these data, it appears that a complex time-dependent program of transcriptional regulation is in play in the morning to upregulate the activity of R5 neurons (Fig 7j). Upregulation of *unc-104, Srpk79D, Syx1a, Rim,* and *nSyb* suggests that R5 neurons are assembling a greater number of mature active zones for neuronal output. Upregulation of *para* and the predicted KChIP *CG5890*, which should increase the voltage-gated conductance of sodium and potassium ions across the membrane, supports the idea that R5 neurons may be primed for greater action potentials in the morning. Upregulation of the two sodium:potassium/calcium antiporters suggests that intracellular calcium levels are elevated in the morning, again consistent with the idea that these neurons are more active in the morning. Significantly elevated levels of six ARGs also support this conclusion. Finally, there is some suggestion that the elevated activity may result in plasticity in the R5 neurons supported by PKA and CREB signaling.

### R5 neurons exhibit time dependent changes in BRP and calcium response to SD

SD/extended wake results in the upregulation of many synaptic proteins (Gilestro et al., 2009). Most notable is the presynaptic scaffolding protein BRP, important for synaptic release(Matkovic et al., 2013), and is upregulated in the R5 neurons following 12 hrs of SD (Liu et al., 2016). KD of *Brp* in R5 neurons decreases rebound response to SD (Huang et al., 2020), suggesting that it is necessary for accumulating and/or communicating homeostatic drive. We hypothesized that differences in the propensity for R5 to induce sleep rebound in the morning/evening may be due to changes in synaptic strength that can be observed by tracking levels of BRP.

To test this idea, we used the synaptic tagging with recombination (STaR) system to selectively express a V5 epitope-tagged BRP in R5 neurons using the FLP/FRT system (Chen et al., 2014) as previously reported (Liu et al., 2016). We examined BRP at ZT1.5 and ZT9.5 with and without SD and found that BRP levels are higher at ZT1.5 than ZT 9.5 (Fig. 8a, b). Interestingly, 2.5 h SD had no effect on BRP intensity at either time point (Fig. 8b). It is possible that BRP changes in response to 2.5 h of SD are not observable, while a longer 12 h deprivation is required to induce sufficient changes for observation(Liu et al., 2016). As reduced BRP expression in the R5 reduces rebound (Huang et al., 2020), it is possible that clock-dependent changes in expression of BRP and associated presynaptic modifications are driving the difference in rebound observed in morning/evening.

**Figure 8:**
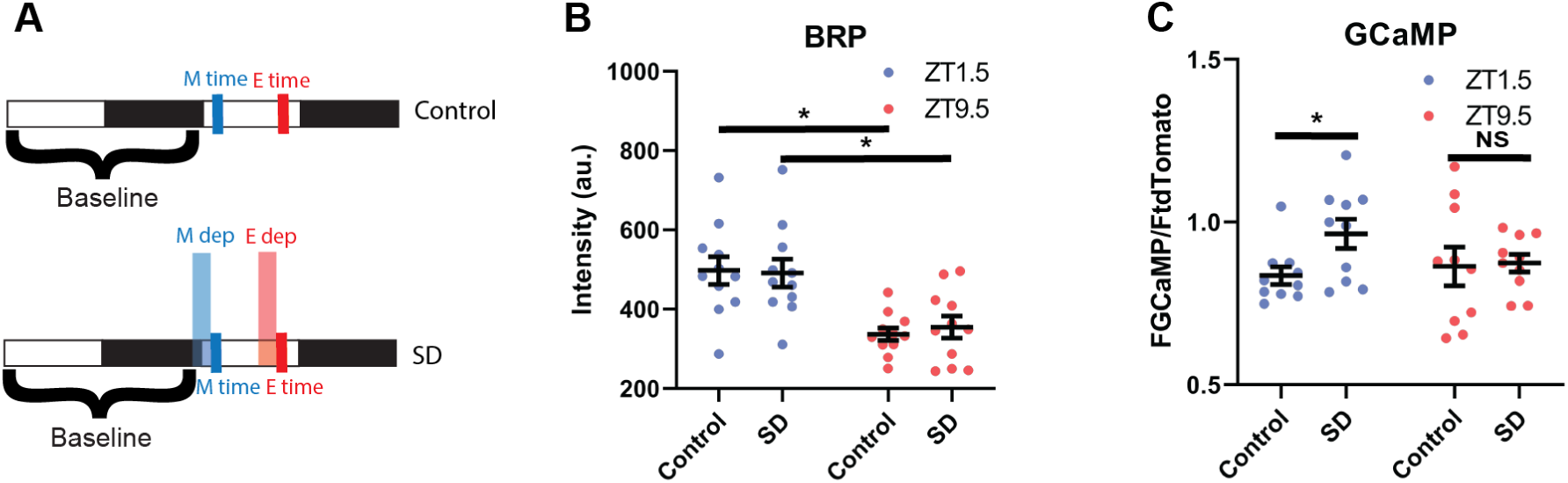
R5 neurons exhibit time dependent changes in BRP and Calcium response to SD. (A) Schematic illustrating deprivation and dissection timing for morning (M) and evening (E) with (lower) and without (upper) SD. (B) Fluorescence of BRP-STaR in R5 projections as a function of time of day and SD. Intensity of BRP staining is decreased in the evening compared to morning in both control (N= 11, 11)(P< 0.001, independent t-test) and SD (N= 11, 11) (P<0.01, independent t-test) groups. Intensity of BRP staining is not affected by SD in the morning (N=11,11) (P> 0.90, independent t-test) or evening (N=11,11) (P>0.58, independent t-test). (C). GCaMP expression in R5 projections (R69F08 > GCamP6s) at ZT1.5 and ZT9.5 with and without SD. GCaMP fluorescence was normalized to the tdTomato fluorescence signal intensity. There is no difference in normalized GCaMP6s signaling between baseline morning (N=10) and evening (N=10) time points. SD in the morning (N=10) increases GCaMP6s intensity (P< 0.05, independent t-test) but not in the evening (N=10) (P>0.87 independent t-test), independent t-test). Data are means +∕- SEM.

The calcium concentration in R5 neurons increases following twelve hours of SD, suggesting that extended wakefulness can induce calcium signaling in these neurons. Blocking the induction of calcium greatly reduces rebound, supporting a critical role for calcium signaling in behavioral output(Liu et al., 2016). Furthermore, R5 neurons display morning and evening cell-dependent peaks in calcium activity across the course of the day indicating that calcium is also modulated by the clock network (Liang et al., 2019). It is unclear whether the circadian clock can modulate wake-dependent changes in calcium activity in the R5 neurons.

To test this idea, we expressed the calcium reporter GCaMP6s (Chen et al., 2013) in the R5 and examined calcium in the morning (ZT1.5) and evening (ZT9.5) with and without SD (Fig. 8a). Interestingly there was no difference between the non-SD flies at each time point (Fig. 8c). This may be because the morning time point resides on the down-swing of the morning-peak of R5 calcium activity while the evening time point resides on the upswing of the evening calcium peak (Liang et al., 2019). Nonetheless, an SD induced increase in calcium was observed in the morning but suppressed in the evening (Fig. 8c), suggesting that the R5 sensitivity to sleep deprivation is gated by the clock.

## Discussion

Here we describe the neural circuit and molecular mechanisms by which discrete populations of the circadian clock network program the R5 sleep homeostat to control the homeostatic response to sleep loss. We developed a novel protocol to administer brief duration SD and robustly measure homeostatic rebound sleep. Using this strategy, we demonstrated that homeostatic rebound is significantly higher in the morning than in the evening. We then identified distinct subsets of the circadian clock network and their downstream neural targets that mediate the enhancement and suppression of morning and evening rebound respectively. Using unbiased transcriptomics, we observed very little gene expression significantly altered in response to our 2.5 h sleep deprivation. On the other hand, we did identify elevated expression of activity-dependent and presynaptic genes in the morning independent of sleep deprivation. Consistent with this finding, we also observe elevated levels of the presynaptic protein BRP. These baseline changes are accompanied by an elevated calcium response to sleep deprivation in the morning mirroring the enhanced behavioral rebound in the morning. Taken together, our data support the model of a circadian regulated homeostat that turns the homeostat up late at night to sustain sleep and down late in the day to sustain wake.

Our studies suggest that homeostatic drive in the R5 neurons is stored post-transcriptionally. As part of our studies, we developed a novel protocol using minimal amounts of SD which could be useful for minimizing mechanical stress effects and isolating underlying molecular processes crucial for sleep homeostasis. 6-24 hours of SD in *Drosophila* is commonly used despite the potential stressful or even lethal effects(R. W. Fernandez et al., 2014; Shaw et al., 2002; Vaccaro et al., 2020). Here we demonstrate that shorter 2.5 hour deprivations not only induce a robust rebound sleep response (Fig. 2), but also the percent of sleep lost recovered at ZT0 is close to 100% versus 14-35% seen in 12 h SD protocols(Blum et al., 2021; Kayser et al., 2014; Nall & Sehgal, 2013; Oh et al., 2014). Using this shorter SD, we now find that many effects observed in R5 neurons with 12 h SD (e.g., increased BRP and upregulation of *nmdar* subunits) are no longer observed with shorter SD, even though the necessity of R5 neurons for rebound is retained after 2.5 h SD (Fig. 6e,f). Previously, translating ribosome affinity purification (TRAP) was used to show upregulation of *nmdar* subunits following 12 h SD(Liu et al., 2016). FACS and TRAP are distinct methodologies for targeted collection of RNA for sequencing and can yield unique gene lists(Cedernaes et al., 2019). One possibility is that upregulation of *nmdar* subunits is occurring locally in neuronal processes, which are often lost during FACS, and/or is at the level of translation initiation or elongation. Nonetheless, in agreement with previous work, we observed SD-induced increases in calcium correlated with behavioral rebound, suggesting that this process is a core feature of the cellular homeostatic response.

Using genetically targeted “loss-of-function” manipulations, we have defined small subsets of circadian clock neurons and downstream circuits that are necessary for intact clock modulation of sleep homeostasis. The use of intersectional approaches enabled highly resolved targeting not possible with traditional lesioning experiments in the SCN(Easton et al., 2004). Collectively our studies defined a Glu^+^ DN1p-TuBu-R4m circuit important for enhancing morning rebound as well as a discrete group of LNds important for suppressing evening rebound. Importantly, most of these effects on sleep rebound are evident in the absence of substantial changes in baseline activity, despite other studies indicating their necessity for normal circadian behavior. Of note, the proposed roles of the DN1p and LNd clock neurons are sleep(Guo et al., 2016) and wake promotion(Guo et al., 2018) consistent with our findings after sleep deprivation. We hypothesize that by using chronic silencing methods, baseline effects may not be evident due to compensatory changes but that these effects are only revealed when the system is challenged by sleep deprivation. Similar genetic strategies in mammals (see (Collins et al., 2020)) may be useful in uncovering which SCN neurons are driving circadian regulation of sleep homeostasis given the comparable suppression of sleep rebound in the evening in humans (Dijk & Czeisler, 1994, 1995; Dijk & Duffy, 1999; Lazar et al., 2015). Nonetheless, the finding of sleep homeostasis phenotypes in the absence of significant baseline effects suggests that a major role of these clock neuron subsets may be to manage homeostatic responses.

Our studies suggest that circadian and homeostatic processes do not compete for influence on a downstream neural target but that the circadian clock programs the homeostat itself. Using an unbiased transcriptomic approach, we discovered time-dependent expression of activity dependent and presynaptic genes (Fig. 7), consistent with previous data that the R5 neurons exhibit time-dependent activity(Liang et al., 2019; Liu et al., 2016). We observed significant upregulation of several genes involved in synaptic transmission (*Syx1a, Rim, nSyb, unc-104, Srpk79D, para, CG5890*) evincing a permissive active state for R5 neurons in the morning. This is accompanied by elevated levels of the key presynaptic protein BRP in the morning compared to evening. It is notable that elevated BRP in the morning is the opposite of what would be expected based on a sleep-dependent reduction in BRP proposed by the synaptic homeostasis hypothesis(Tononi & Cirelli, 2014), suggesting a sleep-wake independent mechanism. Previous studies have shown that modulation of BRP levels in the R5 are important for its sleep function(Huang et al., 2020), suggesting that changes in BRP levels impact R5 function. We hypothesize that these baseline transcriptomic changes underlie the differential R5 sensitivity to sleep deprivation is evident as calcium increases in the morning and not the evening. Indeed, trancriptomic and proteomic studies of the mouse forebrain across time and after sleep deprivation are consistent with the model that the circadian clock programs the transcriptome while homeostatic process function post-trranscriptionally(Bruning et al., 2019; Noya et al., 2019), paralleling what we have found for R5. It will be of great interest to understand the circuit and molecular mechanisms by which circadian clocks regulate the R5 neuronal calcium and synaptic properties and whether similar circuit architectures underlie daily mammalian sleep-wake.

## Acknowledgements

We would like to thank the bloomington stock center and the vienna drosophila resource center for reagents. We thank the Flow Cytometry Core and NU seq at Northwestern University for their assistance in cell sorting and sequencing. We are grateful to the members of our neighbors in the Gallio, Bass and Turek labs for their advice. This work was supported by National Institutes of Health (NIH) grant (R01NS106955), Dept. of Army grant (W911NF1610584), Training Grants in Circadian and Sleep Research (HL 7909-19 and HL007909), and postdoctoral NRSA grant (NS110183).

## Author contributions

R.A., C.R., and T.A.;Methodology, E.O and S.S.; Software, R.A., C.R., and T.A.; Conceptualization, T.A., C.R.; Investigation, T.A., C.R., and S.S.; Formal Analysis, T.A.,C.R., and E.O.; Data Curation, R.A., C.R., and T.A.;Writing-Original Draft, W.K.;Validation, R.A. and C.R.; Supervision, R.A. and C.R.; Project Administration, R.A. and C.R., and T.A.;Funding acquisition.

## Methods

### Fly husbandry and strains

Flies were maintained on a media of sucrose, yeast, molasses, and agar under 12:12 LD cycles at 25°C. 1-3 day old female flies were separated and maintained on standard cornmeal-yeast medium under 12:12 LD cycles at 25°C for 4 nights before experiments began. *Clk*[out] (56754), *per*^s^ (80919), *pdf*-*Gal4* (6899), pBDP (*pBDP-Gal4Uw*)(68384), pBDP split (*p65-AD Uw; Gal4-DBD Uw*) (79603), *R23E10-Gal4* (49032), *R69F08*-*Gal4* (39499), *R58H05 p59AD* (70750), *R48H04 DBD* (69353) *pdf*-*Gal80* (80940), *R51H05 p65AD* (70720), *R18H11 DBD* (69017), *R92H07-Gal4* (40633), *R52B02*-*Gal4* (38814), *R20D01*-*Gal4* (48889), BRPstar (55751), *UAS*-*GCaMP6s* (42746), *UAS*-*TNT* (28838), *UAS*-*kir2.1* (6596) and *UAS*-*hid* (65403) were obtained from the Bloomington Drosophila Stock Center. *mGluR*-RNAi (1793) was obtained from Vienna Drosophila Resource Center. *MB122B* and 2*0xUas-IVS-Syn-GFP* was obtained from Janelia Farm.

### Behavioral assays

Following aging and entrainment, 4-7 day old flies were placed in individual 5×65 mm glass capillary tubes containing sucrose-agar food (5% sucrose and 2% agar). These were then loaded into the Drosophila activity monitor (DAM) system (Trikinetics, Waltham, Massachusetts, USA) and placed in either an empty incubator or, in the case of SD experiments, on a multi-tube vortexer (VWR-2500) fitted with a mounting plate (Trikinetics, Waltham, Massachusetts, USA).

For SD experiments 3 nights (with 2 full days) of undisturbed sleep in 12:12 LD cycling at 25°C served as an acclimation period and baseline. Following the baseline period, SD mechanical stimuli was performed as previously described (Nall & Sehgal, 2013). A 2 second vibration stimulus was applied approximately every 20 seconds with a randomized protocol for a time period of 2.5 hours. In the case of the forced desynchrony protocol this 2.5 hour stimulus was repeated every 7 hours (allowing for a total of 4.5 hours of rest following each stimulus) 24 times until SD occurred at each hour around the clock (Fig. 1a). In abridged experiments this 2.5 hour stimulus was applied 5 times: ZT0, ZT8 and ZT23 of day 3, ZT7 of day 4 and ZT6 of day 5.

For sleep analyses DAM data was processed using custom Java and MATLAB based software. Activity was measured in 1 minute bins and sleep was identified as 5 minutes of inactivity (Hendricks et al., 2000). For SD experiments only flies deprived of >90% of baseline sleep at each SD interval were analyzed (Pfeiffenberger & Allada, 2012). Sleep gain was calculated as the difference between sleep during rebound and sleep during the equivalent 4.5 hours at baseline. Activity actograms were plotted with Counting Macro as previously described (Pfeiffenberger et al., 2010a, 2010b).

### Immunostaining

Following aging and entrainment, 4-7 day old flies were placed in individual tubes containing sucrose-agar food (5% sucrose and 2% agar) for 3 nights. Brains were dissected in PBS (137mM NaCl, 2.7mM KCl, 10mM Na2HPO4 and 1.8mM KH2PO4) and fixed in 3.7% formalin solution (diluted from 37% formalin solution, Sigma-Aldrich) for 30 minutes at 4°C. Brains were washed with 0.3% PBSTx (PBS with 0.3% Triton-X) 5 times (with 15 minute shaking steps at 4°C) before primary antibody incubation. Primary antibodies were diluted in 0.3% PBSTx with 5% normal goat serum and incubation was done at 4°C overnight. Brains were washed for 5 times with 0.3% PBSTx. Secondary antibodies were diluted in 0.3% PBSTx with 5% normal goat serum and brains were incubated at 4°C overnight. Primary antibody used was mouse anti-V5 (1:800 Invitrogen), Secondary antibody used was Alexa 594 anti-mouse (1:800, Invitrogen).

Images were taken using Nikon C2 confocal at 63x magnification and acquired at 1,024 x 1,024 pixels. Analysis of BRP intensity was performed using Fiji/Imagej similarly to previously reported methods (Liu et al., 2016). First max intensity projections were created from confocal stacks of R5 ring projections. The mean intensity of the R5 ring was analyzed by subtracting the average intensity of an adjacent region (background) from the average intensity of the R5 projections.

### Intracellular Ca2+ measurements

Following aging and entrainment, 4-7 day old *R69F08*-*Gal4* > *UAS*-*GCaMP6s*, UAS-CD4-tdTomato flies were placed in individual tubes containing sucrose-agar food (5% sucrose and 2% agar) for 3 nights. Flies were dissected day 4 and imaged in ice-cold control *Drosophila* physiological saline solution (in mM: 101 NaCl, 1 CaCl_2_, 4 MgCl_2_, 3 KCl, 5 glucose, 1.25 NaH_2_PO_4_, and 20.7 NaHCO_3_, pH 7.2, 250 mOsm) (Flourakis et al., 2015). Brains were held ventral side down by a harp slice grid with silica fibers from ALA scientific. GCaMP and TdTomato signal in the R5 ring neuropil was measured immediately (within 5 min) after dissection at ZT1.5 and ZT9.5. Imaging experiments were performed on an Ultima two-photon laser scanning microscope (Bruker, former Prairie Technologies, Middleton, WI). Images were acquired with an upright Zeiss Axiovert microscope with a 40×0.9 numerical aperture water immersion objective at 512 pixels × 512 pixels resolution. Single optical R5 section was selected and recorded as previously described (Liu et al., 2016). In brief a single optical section was selected based on visual assessment of maximum area of tdtomato signal. The GCaMP signal was recorded at ∼1 fps for 60 seconds. The average projection of the frames was used to calculate the GCaMP and TdTomato signal.

### Connectome analysis

We accessed the NeuPrint API via R using a Natverse-based software package, *neuprintr*, along with two other open-source data visualization tools, *hemibrainr* and *ggplot2 (Bates et al., 2020)*. R scripts provided by the Natverse creators were modified to generate connectivity graphs (node networks) and neuron skeletonizations (visualizations of neuronal morphology). Our modified scripts can be found at https://rpubs.com/eogunlana0827/modified-code-for-analysis. Most of the neurons used in this study were identified based on their annotation in Neuprint. Cry-positive LNds were identified in the total LNd based on morphology according to the images in Schubert et al (Schubert et al., 2018).

To generate node networks for sleep pathways, the body IDs of the pre- and post-synaptic targets were determined by querying the neuron types and storing the retrieved data into two dataframes (A and B, respectively). Once A and B were determined, the shortest paths between the two types were then calculated. The code accounts for any duplicates that may arise when running *neuprintr*’s “shortest paths”function. This information is stored in another dataframe that represents each pre- and post-synaptic neuron instance in the pathway, along with their names/types and the number of synapses between each neuron. Before establishing the network environment in which the data are plotted, the newly created dataframe was modified so that only the pre- and post-synaptic neuron types and synaptic weights were included, thereby removing any body ID information. We then utilized the *network* and *ggnetwork* packages (both under the *ggplot2* package framework) to create the network environment. Colors were assigned to each neuron type using a list of variables provided in the pre-made R scripts. Finally, the connectivity graphs were plotted using *ggplot2* and exported to PDFs.

The *hemibrainr* package was used to generate visualizations of neuronal morphology from the EM data underlying Neuprint *(Bates et al., 2020)*. For each neuron type in the sleep pathways, we collected the neuron mesh data from their NeuPrint body IDs using a hemibrainr function and then stored them in a variable. Then, we randomly sampled a color to assign to each neuron type using a built-in R function. The neuron mesh was then plotted in a 3D environment, and then oriented so that the anterior side of the brain was facing the viewer.

### Fluorescence Activated Cell Sorting and RNA-seq

FACS/RNA-seq was performed as previously reported (Xu et al., 2019). Briefly, flies were housed in DAM system behavior boards in either control or sleep deprivation conditions. Immediately following SD, the boards were recovered from the incubators and transferred to CO_2_ pads. Brains were dissected in ice-cold modified dissecting saline (9.9 mM HEPES-KOH buffer, 137 mM NaCl, 5.4 mM KCl, 0.17 mM NaH_2_PO_4_, 0.22 mM KH_2_PO_4_, 3.3 mM glucose, 43.8 mM sucrose, pH 7.4) with 0.1 μM tetrodotoxin (TTX), 50 μM D(–)-2-amino-5-phosphonopentanoic acid (AP-5), and 20 μM 6,7-dinitroquinoxaline-2,3-dione (DNQX) to block neuronal activity. Following dissection, brains were transferred to SM^Active^ medium (4.18 mM KH_2_PO_4_, 1.05 mM CaCl_2_, 0.7 mM MgSO_4_·7H_2_O, 116 mM NaCl, 8mM NaHCO_3_, 2 mg/ml glucose, 2 mg/ml trehalose, 0.35 mg/ml α-ketoglutaric acid, 0.06 mg/ml fumaric acid, 0.6 mg/ml malic acid, 0.06 mg/ml succinic acid, 2 mg/ml yeast extract with 20% non heat-inactivated FBS, 2 mg/ml insulin and 5mM pH6.8 Bis-Tris) with 0.1 μM TTX, 50 μM AP-5, and 20 μM DNQX on ice while the rest of the brains were dissected. 40-45 brains per time point were pooled as a single sample and every condition and time point was run in triplicate for a total of twelve samples. Following dissection, the brains were pelleted by centrifugation (2000 rpm, 1 min) and washed twice with 500 uL of chilled dissecting saline (containing TTX, AP-5, and DNQX). Dissecting saline was removed and the brains were incubated at room temperature in 100 μL of papain (50 unit/mL, heat activated for 10 min at 37°C) for 30 minutes. Following digestion, the papain was inactivated with 500 μL of chilled SM^Active^ medium and then washed twice with chilled medium on ice. The brains were triturated by pipetting with a flame-rounded 1,000 μL pipette tip (30 times with a medium opening, 30 times with a small opening). The sample was filtered using a 100 μm nylon filter (Sefar Nitex 03-100/32) then transferred to the Northwestern FACS core on ice. GFP-positive cells were sorted on an Aria II FACS Cell Sorter into an extraction buffer from the Arcturus PicoPure Kit. We collected 300-550 cells per sample. Following sorting, the cells were lysed in extraction buffer by incubating at 42°C for 30 min. After lysing, the cells were stored in a -80°C freezer until libraries could be made.

Total RNA was extracted from collected cells using the PicoPure Kit with on-column DNAse I digestion according to manufacturer instructions. Following extraction, the RNA was immediately concentrated down to 1 μL using a Speed-Vac. First strand cDNA was prepared using a T7-oligo-dT primer and SuperScript III following manufacturer instructions. Second strand synthesis was performed with DNA Polymerase (18010025), Second Strand Buffer (Cat#10812014), 10 mM dNTP (18427088), DNA Ligase (18052019), and RNaseH (18021071). The cDNA was used as a template for one round of *in vitro* transcription (IVT) using T7 RNA polymerase and the Ambion MegaScript kit according to manufacturer instructions. IVT was carried out at 37.5°C for 4 hours. Following IVT, the new RNA was purified using a Qiagen RNEasy kit and then used to generate libraries for RNA-seq using an Illumina TruSeq Stranded Kit. Libraries were checked for appropriate size distribution and purity by Bioanalyzer, then sent to Novogene for sequencing. We generated 30 million reads per sample.

Reads were pseudo aligned and quantified using Kallisto (v0.46.1) (Bray et al., 2016) against a prebuilt index file constructed from Ensembl reference transcriptomes (v96). Kallisto was used to process paired end reads with 10 bootstraps. Differential expression analysis of the resulting abundance estimate data was then performed with Sleuth (v0.30.0)(Pimentel et al., 2017). Gene-level abundance estimates were computed by summing transcripts per million (TPM) estimates for transcripts for each gene. To measure the effect of a particular condition against another condition for a variable, sleuth uses a Wald test which generates *p* values as well as *q* values (an adjusted *p* value using the Benjamini-Hochberg procedure).

### Statistics

Statistical analyses and figures were produced with Excel, Matlab and Prism. Paired student T-tests were used to compare 2 groups/time points. Repeated one and two factor ANOVA analyses were used to compare multiple time points/groups with Tukey’s post hoc test. Additional details regarding tests and significance values are provided in the figure legends.

**Supplementary Video 1: Flies exhibit sleep following 2.5 hours SD terminating at ZT1.5** Sped up video recording of 4.5 hours of rebound of 36 WT flies following SD from ZT23-ZT1.5. Hours post SD are indicated in red in the bottom right corner. Flies exhibit little movement throughout the 4.5 hours following SD indicating sleep.

**Supplementary Video 2: Flies are active following 2.5 hours SD terminating at ZT9.5** Sped up video recording of 4.5 hours of rebound of 36 WT flies following SD from ZT7-ZT9.5. Hours post SD are indicated in red in the bottom right corner. After a brief period of immobility flies exhibit high activity (low sleep) preceding lights on.

---

**Supplemental Figure 1:**
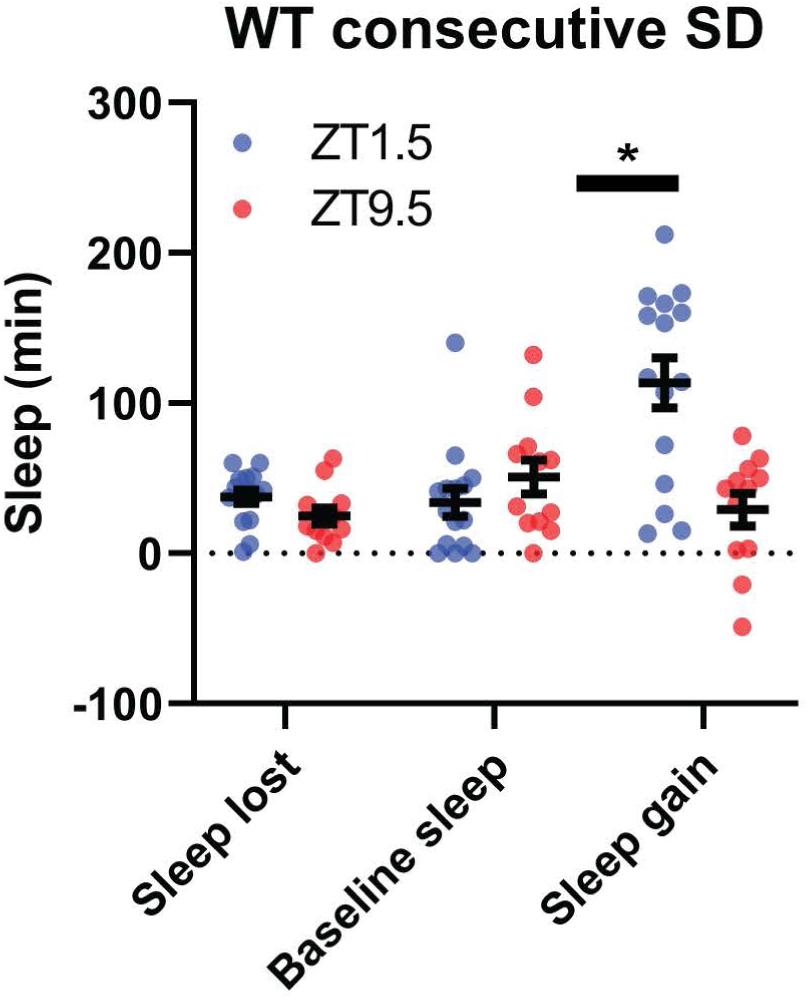
WT flies exhibit higher rebound in the morning than the evening even using abridged protocol. Comparison of sleep lost, baseline sleep, and sleep gain at moming(ZTl .5) and evening (ZT9.5) time points using abridged protocol with WT flies. Sleep gain is greater for WT (N=18) at ZTl.5 compared to ZT9.5 (P<.001, paired t-test).

**Supplemental Figure 2:**
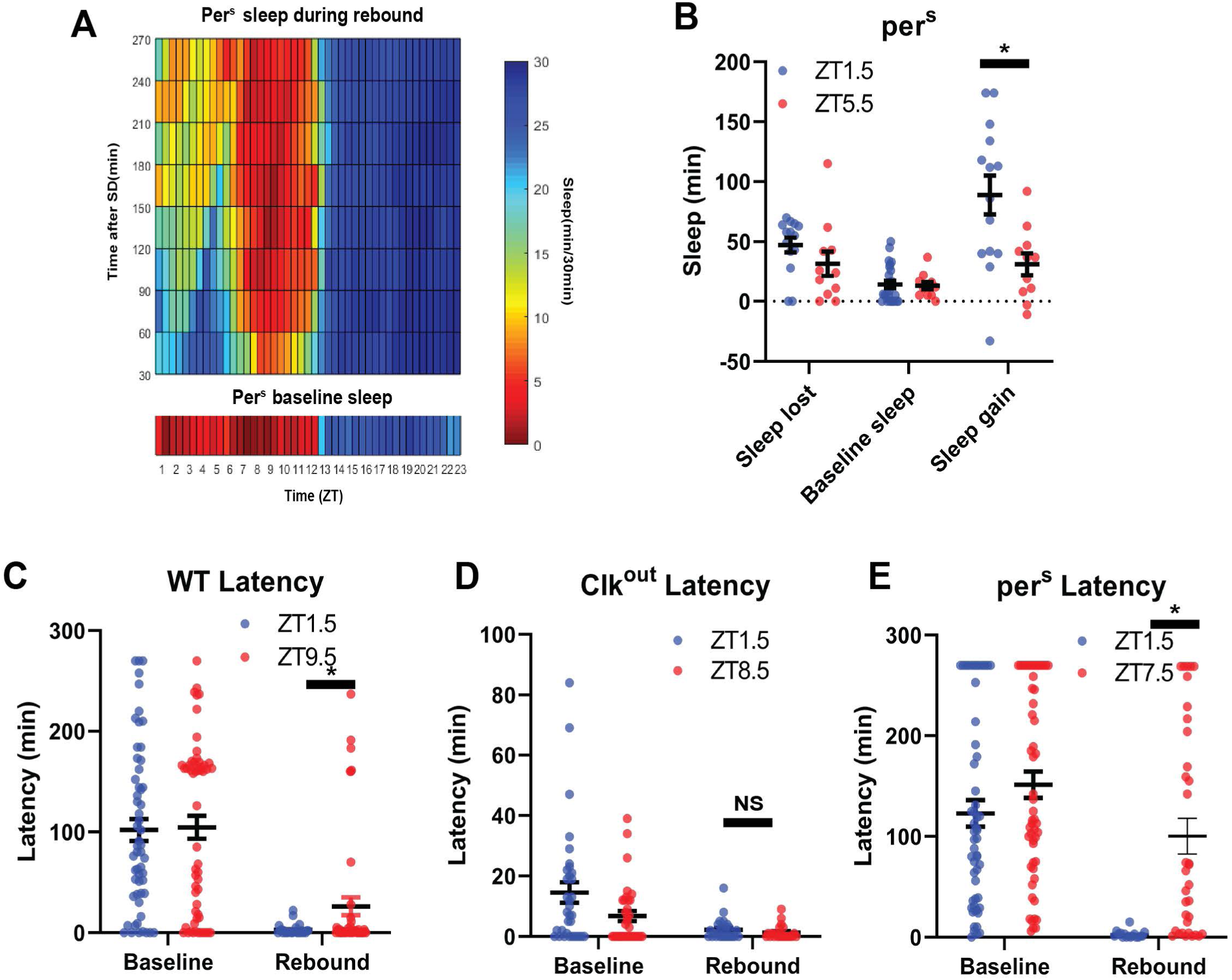
Total sleep and sleep latency vary as a function of time and SD. (A) Rebound sleep heatmap (above) illustrates average sleep as a function of time of day when rebound occurred (ZT) and minutes after FD episode. Missing time points are filled using matlab linear interpolation function. Baseline sleep heatmaps (below) illustrate average sleep during 30 min bins. pers (N=45) baseline sleep displays low sleep following lights on and preceding lights off. Immediately following SD flies show increased sleep except in the hours preceding lights off. Flies tend to sleep less as rebound time proceeds. (B) FD sleep during two baseline time periods (sleep lost and baseline sleep) and sleep gain for rebound occuring at ZTl.5 and ZT5.5. Sleep gain is greater for pers (N=45) rebound at ZTl.5 compared to ZT5.5 (P<.01, paired t-test). (C,D,E) Morning and evening sleep latency at baseline and following deprivation for WT and circadian mutant flies. Morning times are matched with evening time points with similar baseline latency. (A) Following SD, WT (N=32) sleep latency is greater in the evening compared to matched morning time point(P<.05, paired t-test). (B) No difference in sleep latency following SD between matched morning/evening time points in Clkout (N=40) (P>0.50, paired test). (D) Following SD, pers (N=45) sleep latency is greater in the evening compared to matched morning time point (P<.0001, paired t-test).

**Supplemental Figure 3:**
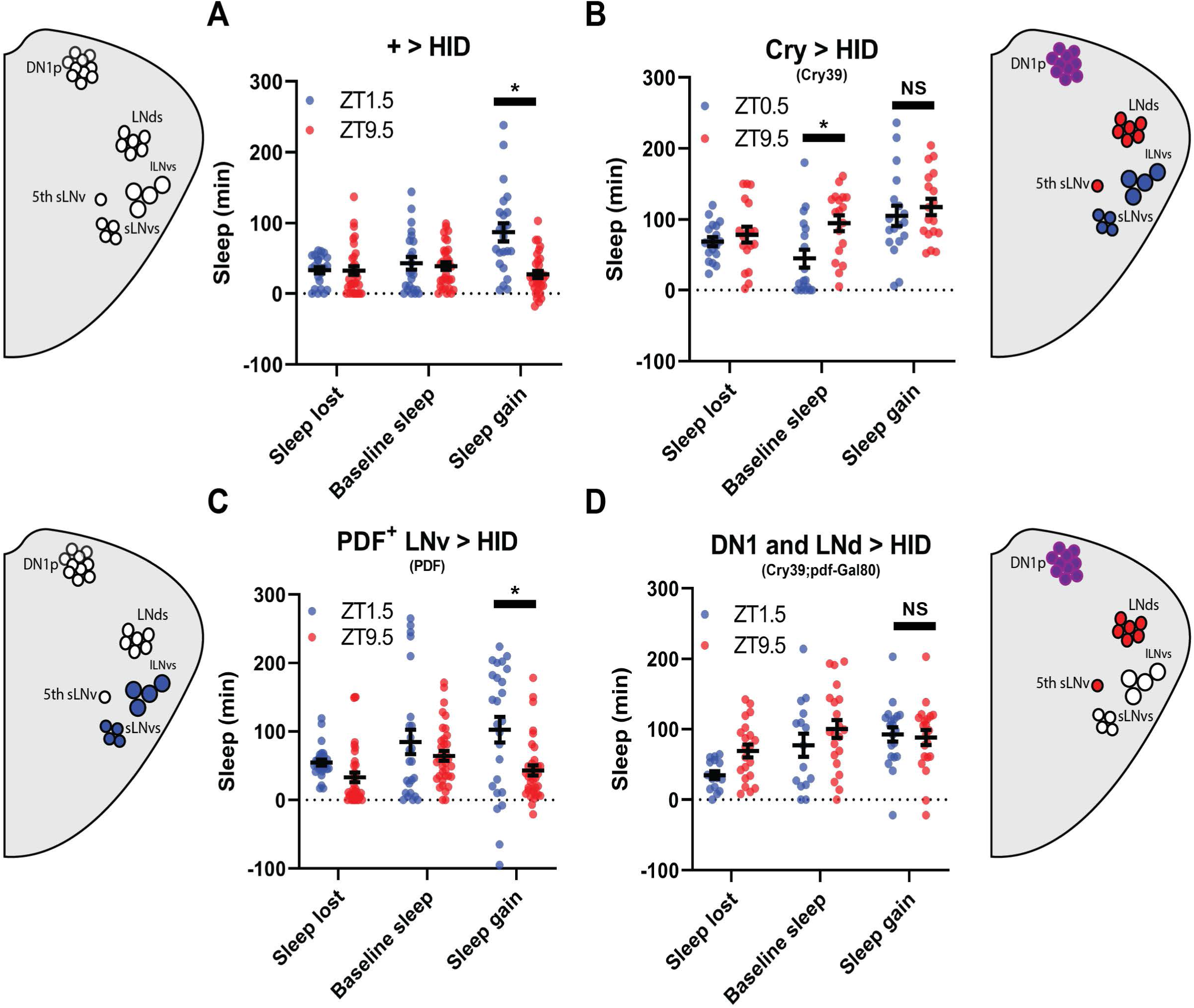
PDF+ neurons do not mediate morning/evening differences in rebound. (A,B,C,D) Comparison of sleep lost, baseline sleep, and sleep gain following deprivation at morning and evening timepoints in clock neuron-ablated flies. Morning times are matched with evening time points with similar baselines. (A) Control flies with no ablated neurons (+ > hid) (N=27) exhibit greater rebound in the morning compared to matched evening time point (P<0.0001, paired t-test). (B) Flies with most clock neurons ablated (cry39 > hid) (N=19) exhibit no difference in sleep gain between matched morning/evening time points (P>0.70, paired t-test). (C) Files with PDF+ neurons ablated (pdf > hid) (N=35) exhibit greater rebound in the morning compared to a matched evening time point (P<0.01, paired t-test). (D) Flies with most clock neurons ablated except PDF+neurons (cry39; pdf-Gal80 > hid) (N=22) exhibit no significant difference in sleep gain between matched morning/evening time points (P>0.97, paired t-test).

**Supplemental Figure 4:**
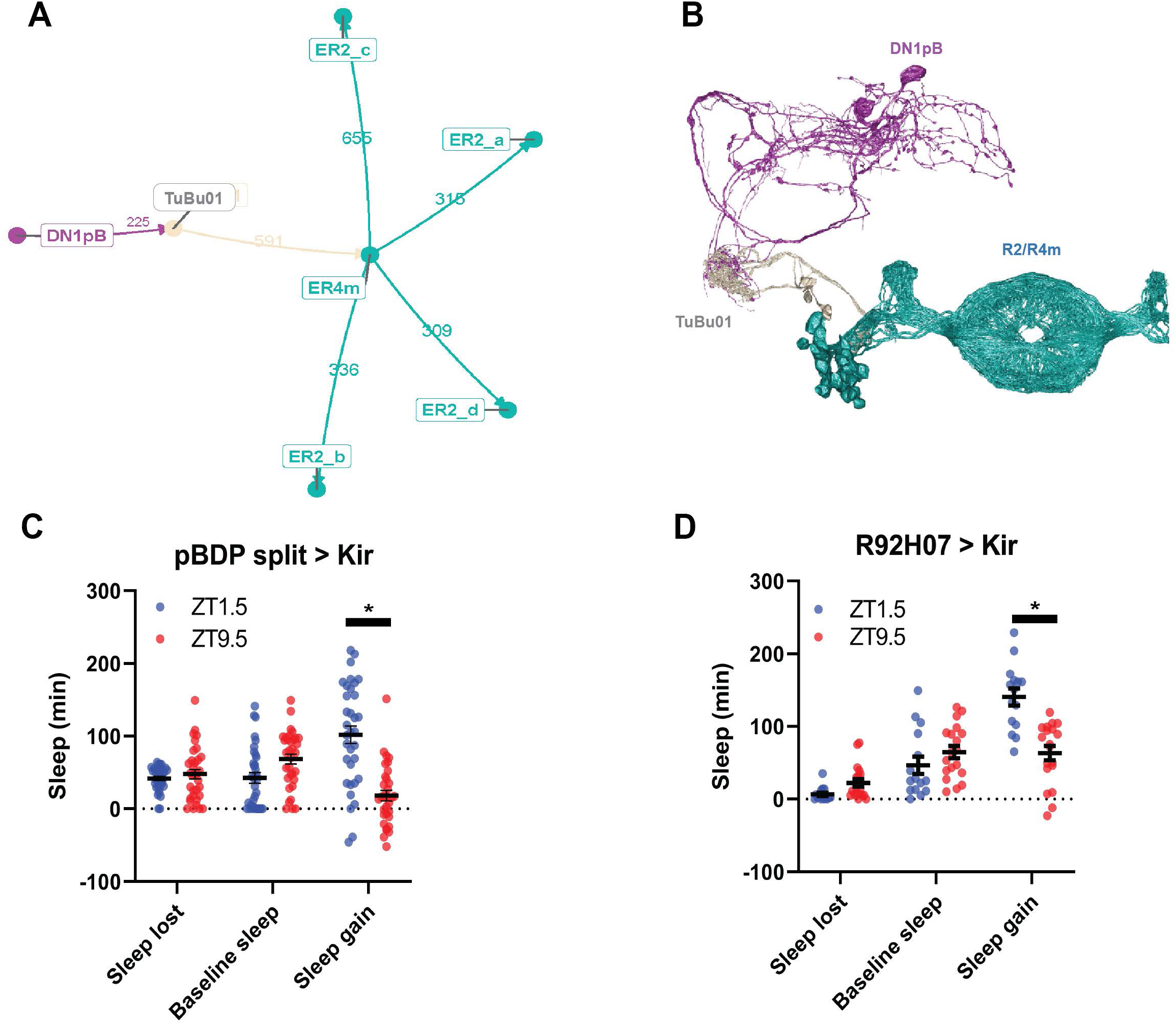
Connectome analysis demonstrates link between anterior projecting DNlps and R2/R4m ellipsoid body ring neurons. Node network diagram of pathway from anterior projecting DNlps to R2/R4m via Tubu intermediates. Arrows indicate directionality of projections and numbers represent average synaptic connections between groups of neurons. Dorsal view of neuronal morphology of pathway from anterior projecting DNlps to R2/R4m via Tubu intermediates according to Neuprint EM reconstruction. Each cell subtype is color coded to match the model in A. (C-D) Comparison of sleep lost, baseline sleep, and sleep gain following deprivation at morning and evening time­ points while modulating TuBu neurons. Morning times are matched with evening time points with similar baselines. Enhancerless-Gal4 control flies (pBDP > Kir) (N=2 **l)** exhibit greater rebound in the morning compared to a matched evening time point (P<0.01, paired t-test).(D) Flies with TuBu neurons silenced (R92H07 > Kir) (N=21) exhibit greater rebound in the morning compared to a matched evening time point (P<0.0001, paired t-test).

**Supplemental Figure 5:**
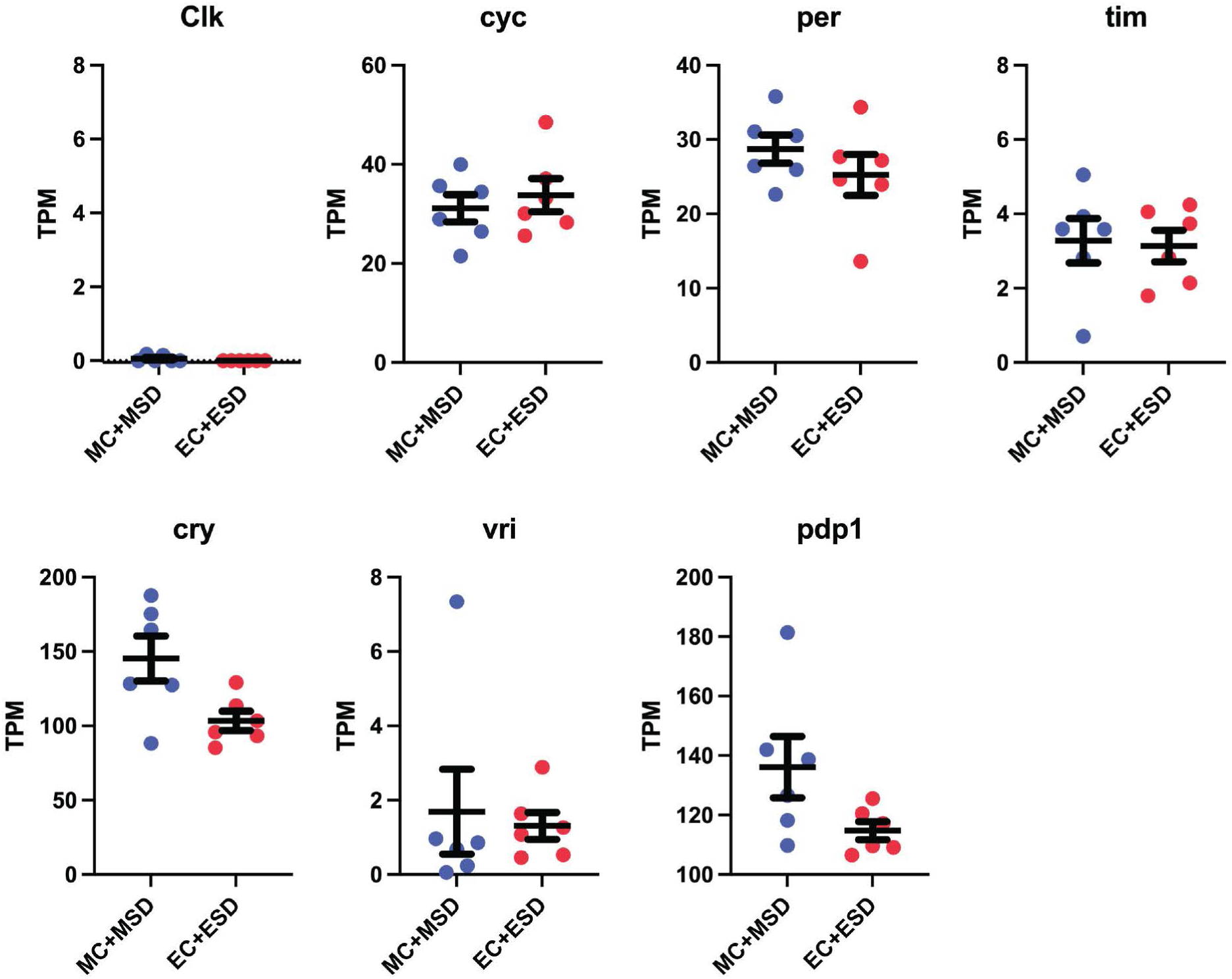
Clock genes are not oscillating in RS neurons. Scatter plots for core clock genes. Transcripts Per Kilobase Million (TPM) is shown for each sample. All morning samples are grouped and all evening samples are grouped.

**Supplemental Table 1:** Summary of male morning and evening anticipation. Data are means +/- SEM (*p<0.05, **p<0.01, ***:p<0.001).

| Genotype | Region/Cells targeted | LD morning anticipation | LD evening anticipation | N |
| --- | --- | --- | --- | --- |
| **+> hid** | clock Gal4 control (HID) | 0.14 +/- 0.04 | 0.37 +/- 0.03 | 17 |
| ***pBDP split > hid*** | Split control (HID) | 0.13 +/- 0.02 | 0.24 +/- 0.03 | 26 |
| ***pBDP split > kir*** | Split control (Kir) | 0.10 +/- 0.02 | 0.33 +/- 0.04 | 12 |
| ***cry39 > hid*** | broad clock | 0.05 +/- 0.02 ** | 0.12 +/- 0.04 *** | 30 |
| ***pdf > hid*** | PDF | -0.07 +/- 0.02 *** | 0.24 +/- 0.03* | 38 |
| ***cry39; pdf-gal80 > hid*** | LNd and Dn1 | 0.05 +/- 0.01 * | 0.06 +/- 0.04 *** | 14 |
| ***R51H05 AD ; R18H11 DBD > hid*** | Glu+ DN1p | 0.06 +/- 0.02 * | 0.25 +/- 0.02 | 22 |
| ***MB122 > hid*** | 3-4 LNds PDF-sLNv | 0.12 +/- 0.02 | 0.25 +/- 0.02 | 35 |
| ***MB122 > kir*** | 3-4 LNds PDF-sLNv | 0.07 +/- 0.02 | 0.22 +/- 0.02 | 26 |

